# Structural basis of the lobster carapace blue colour mediated by an HPR protein

**DOI:** 10.64898/2026.02.26.708136

**Authors:** Maria Claudia Cedri, Harsh Bansia, Adolfo Amici, Maria Grazia Ortore, Andrew McCarthy, Christoph Mueller-Dieckmann, Rafael Lingas, Bo Durbeej, Nadia Raffaelli, Tong Wang, Amedee des Georges, Michele Cianci

**Author notes:** Department of Molecular Pathobiology, NYU College of Dentistry, 433 1st Avenue, New York, NY 10010, USA.

## Abstract

The chemical basis underlying the striking blue hue of live *H. americanus,* known as American lobster, are studied in evolutionary biology and in polyene physical chemistry. Carapace colouration is generated by the antioxidant astaxanthin bound within the carotenoprotein crustacyanin complexes. Here, we present the ex vivo structure of the most abundant α-crustacyanin and β-crustacyanin forms, determined respectively by cryo-electron microscopy and X-ray crystallography to a resolution of 2.75 Å. Our structural analysis reveals α-crustacyanin as an elongated arrangement of β-crustacyanin heterodimers tethered by an heptatricopeptide repeat (HPR) protein. In vitro complex formation between the β-crustacyanin unit with a synthetic heptatricopeptide reproduces the observed blue colour of α-crustacyanin, identifying the HPR protein, in concert with crustacyanins, as contributor in tuning carapace colour. Overall, these results explain how nature adjusts the colour across the entire visible spectrum by exploiting the bathochromic shift of astaxanthin from its unbound red form (λ_max_ = 472 nm) firstly to the β-crustacyanin violet bound form (λ_max_ = 591 nm), and then to the α-crustacyanin bound blue form (λ_max_ = 631 nm).

## Introduction

Nature’s canvas is adorned with a mesmerizing array of hues, each shade serving as a testament to the intricacies of evolutionary adaptation. From vibrant plant pigments to the iridescent sheen of animal coats, natural colouration stands as a masterpiece of survival strategy. It serves as both shield and signal, warding off predators, protecting against ultraviolet damage, and conveying vitality and desirability during the mating season. Within this intricate tapestry of survival strategies, the study of natural colouration unveils profound insights into adaptation and communication mechanisms ingrained in the biological world (Cuthill et al., 2017). Plants get their variety of colours from pigments like chlorophyll, carotenoids, betalains and anthocyanins, while animals use melanin, tetrapyrroles, or carotenoids sourced from feed. Marine invertebrates display a wide-range of colours, with notable examples including the striking blue of the starfish *Linckia laevigata* (Echinodermata: Asteroidea), and the ethereal blue tones of the jellyfish *Velella velella*, commonly known as by-the-wind sailor. *H. americanus*, known as American lobster, and *Homarus gammarus* or European lobster (Holthuis, 1991) are closely related and widely recognized for the deep slate-blue hue when alive. The blue pigments found in the starfish *Linckia laevigata* (S. T. Williams, 2000; Zagalsky et al., 1989), the jellyfish *Velella velella* (Zagalsky PF & Herring PJ, 1977) and the lobster *Homarus gammarus* (Cheeseman et al., 1966; Zagalsky & Cheeseman, 1963) and *Homarus americanus* (Zagalsky & Tidmarsh, 1985), along with several other marine species (Milicua et al., 1986; Wade et al., 2009) have been isolated and characterised as carotenoproteins. Carotenoproteins are large protein-carotenoid complexes, common throughout the invertebrates, responsible for the colouration of the external surface of the animal and internal tissues, like the blood, eggs, ovary, hypodermis, and stomach wall (Zagalsky, 1985). The blue colouration of the lobster carapace is generated by the exocuticle carotenoproteins α-, β-, and γ-crustacyanins (Cheeseman et al., 1966; Quarmby et al., 1977). β- and α-crustacyanins alone represent about 80-90 % of carotenoproteins present in lobster carapace (Zagalsky & Tidmarsh, 1985). About 10-20% of the carapace carotenoprotein is as γ-crustacyanin, distinct from α- and β-crustacyanin but identical in amino acid composition (Zagalsky, 1985).

α-crustacyanins are constituted by multiple copies of the β-crustacyanin heterodimer in which the chromophore, namely the carotenoid astaxanthin (3,3’-dihydroxy-β,β-carotene-4,4’-dione), is associated stoichiometrically 1:1 with the protein subunits CRTC and CRTA (Zagalsky, 1985). The absorption spectrua λ_max_ of astaxanthin moves from λ_max_= 472 nm in the unbound form, producing a visible red colour, to a λ_max_= 591 nm in the β-crustacyanin form (Cheeseman et al., 1966; Zagalsky & Cheeseman, 1963; Zagalsky & Tidmarsh, 1985) generating a blue-violet colour, to a λ_max_= 625 nm in the γ-crustacyanin form and finally to λ_max_= 631 nm in α-crustacyanin form.

Astaxanthin spans then the entire visible spectrum in its unbound and bound states, presenting opportunities in the food and nutraceutical industries as a versatile food colorant with added antioxidant benefits. Notably, the α-crustacyanin pigment has a shade and brilliance similar to an artificial colourant, the Federal Food, Drug, and Cosmetic Act (FD&C) Blue No. 1, otherwise known as Brilliant Blue No. 1, which has a peak of absorption of the visible light at 629 nm (Newsome et al., 2014). Astaxanthin also has a strong quenching effect against singlet oxygen, with antioxidant activities approximately ten times stronger than those of β−carotene, and hundred times greater than those of a tocopherol (Miki, 1991). Astaxanthin is also more effective than β-carotene and lutein at preventing UV light photooxidation of lipids (Santocono et al., 2006). Recently, it has been shown that ultrafast singlet exciton fission (Smith & Michl, 2010) occurs in astaxanthin aggregates (Musser et al., 2015), making them a case study to unravel the full potential of singlet fission in solar energy harvesting (Musser et al., 2015).

The structure of the blue-violet coloured β-crustacyanin from *H. gammarus* elucidates the basis for the bathochromic shift of the carotenoid spectra to λ_max_ = 591 (Cianci et al., 2002). β- crustacyanin is a heterodimer of two lipocalin proteins with molecular weights of 21 kDa (CRTC) (Cianci et al., 2001; Gordon et al., 2001; Habash et al., 2004) and 19 kDa (CRTA) and two non-covalently bound astaxanthin molecules (Cianci et al., 2002).

Investigations with samples purified from *H. gammarus*, by small angle X-ray scattering (SAXS) (Chayen et al., 2003; Dellisanti et al., 2003) and negative stain electron microscopy (Rhys et al., 2011) postulated various possible arrangements of β-crustacyanins to form α-crustacyanin up to a resolution of around 30 Å suggesting that an integrative structural approach could provide additional information.

Crustacyanin apo-proteins from *H. americanus* H1 and H2 are closely related to the ones of *H. gammarus* (CRTC and CRTA) with sequence identity higher than 96% (Cedri et al., 2026; Ferrari et al., 2012). Here, through a combination of X-ray crystallography, cryo-electron microscopy (cryo-EM) and SAXS, we report the 2.75 Å resolution structures of β-crustacyanin and α-crustacyanin, both purified from ex vivo material from *H. americanus*.

The cryo-EM structure of *H. americanus* α-crustacyanin complex reveals, besides the expected β-crustacyanin units, the presence of a third protein, a heptatricopeptide repeat protein (HPR). This HPR protein presents a coil-helix-coil fold and wraps around the entire complex. It binds a β-crustacyanin subunit at each repeat, interacting with it at the carotenoid binding region, and stabilizing the pseudo helical oligomeric assembly. The analysis of the atomic resolution structures of the crustacyanin complexes identify contributors to the bathochromic shift of the carotenoid spectrum from λ_max_ = 591 nm to λ_max_ = 631 nm. In vitro complex formation between the purified β-crustacyanin unit and a synthetic heptatricopeptide reproduces the observed blue colour of α-crustacyanin, identifying HPR protein, in concert with crustacyanins, as key contributor in tuning carapace colour.

## Results

### Structure of *H. americanus* β-crustacyanin heterodimer structure and comparison with homolog from *H. gammarus*

The purified β-crustacyanin absorbed at λ_max_ = 591 nm, and it yielded crystals diffracting to 2.75 Å resolution (Cedri et al., 2026). The crystal structure of β-crustacyanin heterodimer from *H. americanus* is consistent with the *H. gammarus* β-crustacyanin crystal structure already determined (Cianci et al., 2002). The overall fold of the heterodimer is fully conserved (Fig. S1) with the *H. gammarus* counterpart, as expected from the 96.7% and 98.3% sequence identity for the H1 and H2 subunits from *H. americanus* respectively (Cedri et al., 2026). The availability of diffraction data from *H. americanus* β-crustacyanin crystal at 2.75 Å resolution allowed to confirm a network of water molecules in proximity of the astaxanthin end ring binding sites (Fig. S2), which were only partially observed in the β-crustacyanin heterodimer from *H. gammarus* at 3.2 Å resolution. The presence of water molecules in the surrounding of the astaxanthins had long been postulated before the availability of the first crystal structure of β-crustacyanin, and it is considered essential to favour the polarization of astaxanthin by the surrounding amino acid side chains (Britton et al., 1997).

The protein sequence alignment of the subunits H1 and H2 from *H. americanus* versus C1 and A2 from *H. gammarus* reveals only a limited number of point mutations (Cedri et al., 2026). In H1 the mutations are N5D, S30N, K61E and K66T, while in H2 the mutations are H48N, T55G and T147C. The T147C mutation generates an additional disulfide bridge between Cys115 – Cys147 in the H2 subunit, whereby for the A3 subunit of the β-crustacyanin heterodimer from *H. gammarus*, only two disulfide bridges have previously reported (Cianci et al., 2002) (Fig. S1B).

All the mutations are solvent exposed amino acids, away from the astaxanthin binding pockets, where the main difference that occurs is a 5 Å shift of the Phe101(H1) phenyl ring towards a hydrophobic void between the two astaxanthin molecules. In the *H. gammarus* β-crustacyanin, the Phe101(H1) phenyl ring was clearly positioned above the astaxanthin molecules, and a spurious electron density was observed above and assigned as a dodecane molecule, possibly coming from the paraffin oil used in the batch under oil crystallization conditions (Cianci et al., 2002). In the *H. americanus* β-crustacyanin structure there is no evidence of any additional electron density, which agrees with the fact that crystals of *H. americanus* β-crustacyanin have been obtained with the standard hanging drop set up. The different position of Phe101 closer to the two astaxanthin molecules is not influencing the absorption spectra of *H. americanus* β-crustacyanin, which was measured at 591 nm, in line with one of *gammarus* β-crustacyanin, reported to be in the region of 585-591 nm (Zagalsky, 1985).

The two lipocalin subunits H1 and H2 interact via the loop region connecting strand G and H (Cianci et al., 2002) of each subunit and bind two astaxanthin molecules (Fig. S1A). The two bound astaxanthins have a *6-s-trans* conformation with the end rings essentially coplanar with their polyene chains. The carotenoids approach within 6.7 Å at the central portions of their respective polyene chains. For each astaxanthin molecule we define the C1-6 end ring as the one located within the calyx of the lipocalin and the C1’-6’ end ring located at the interface region of two lipocalin subunits H1 and H2 (Fig. S2). The end ring C1-6 of AXT1 is within hydrogen bond distance of the H1 subunit residues Gln46 and Asn54, and constrained by Phe86, Ile95 and Phe134 (Fig. S2A). At the C1-6 end ring of the astaxanthin, positioned within the H1 lipocalin calyx, two water molecules are present (W1 and W2), tethered by a network of hydrogen bonds between the astaxanthin and the surrounding amino acids (Tyr56 and Tyr97). The end ring C1’-6’ is located in proximity of Phe86, His90, Pro102, of the H2 subunit, and Ile3, Phe126, Tyr128 of the H1 subunit (Fig. S2C).

Conversely, for the AXT2 molecule, the end ring C1-6 is in proximity of the H2 subunit residues Gln32, Gln41, Ser49, Tyr51, Ile76, and Phe133 (Fig. S2D). One water molecule (W3) is found here, tethered by a network of hydrogen bonds between the carbonylic oxygen of astaxanthin and the side chain of Ser49, Tyr51 and Thr64. The end ring C1’-6’ is in proximity of Asn86, His92, Ser94, and Pro104 of H1 subunit, and Ile3, Asp123 and Tyr125 of H2 subunit (Fig. S2B). A water molecule (W4) is also found here near the hydroxy group of C1’-6’ end ring (Fig. S2B). Two out of four water molecules observed in our studies are consistent with the previously reported structure of the *H. gammarus* β-crustacyanin (Cianci et al., 2002). The other two have not been observed before. Both astaxanthin polyene chains, C7 to C7’, are clamped in a hydrophobic patch of Phe, Tyr, Val and Pro residues, which locks the position of the central methyl groups C20 and C20’. The protein environment of the two end rings of both astaxanthin in both *H. gammarus* and *H. americanus* is therefore conserved and similarly asymmetric with a partial positive charge at the end ring C1-6, and a partial negative charge at the end ring C1’-6’.

### Structure of *H. americanus* α-crustacyanin presents a pseudo helical architecture

Purified *H. americanus* α-crustacyanin, yielded diffracting crystals albeit only to a resolution of 6.3 Å (Cedri et al., 2026). This resolution was insufficient to solve the crystal structure by molecular replacement using multiple copies of the β-crustacyanin heterodimer reported here. Initial negative stain data confirmed the monodispersity of the α-crustacyanin particles (Cedri et al., 2026). This led us to attempt to solve the structure by single-particle cryo-EM. Imported particles underwent iterative rounds of reference-free 2D classification (Fig. 1A) to select the 369,648 particles for 3D reconstruction (Fig. 1B). We performed two further 3D reconstruction protocols. A homogeneous 3D refinement of the 369,648 particles yielded a map at the resolution of 2.86 Å interpreted as a pseudo helical particle, of approximately 281 Å in length and formed by oligomerization of apparently twelve β-crustacyanin subunits.

**Fig. 1.**
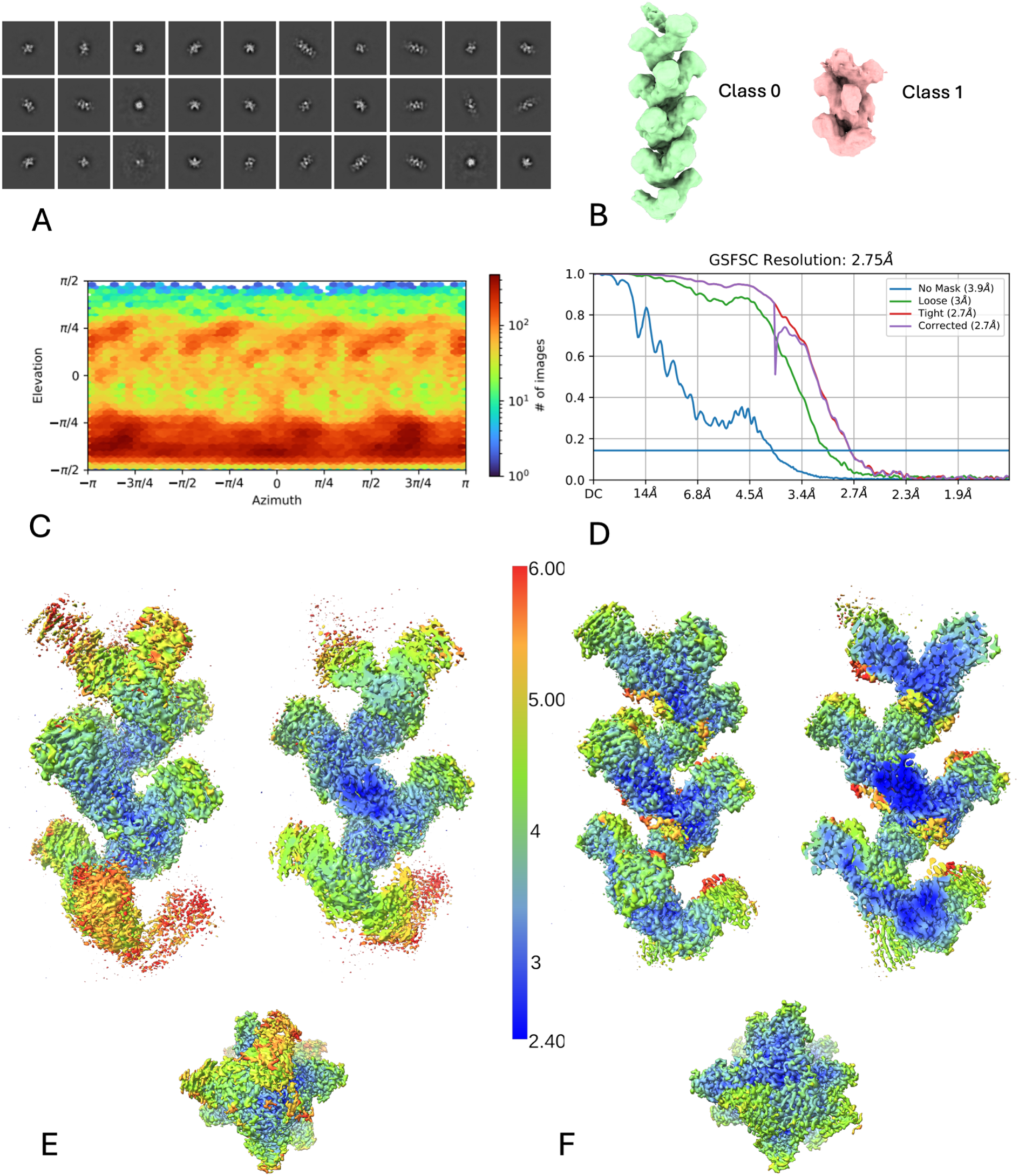
Structure solution diagram of cryo-EM reconstruction of ex vivo α-crustacyanin complex from *H. americanus*. **A** representative 2D classes. **B** volumes of class 0 and 1 obtained from heterogeneous refinement. **C** orientation plots contain information regarding the distribution of orientations for the final datasets. **D** FSC (Fourier Shell Correlation) plot. The resolution is reported at the gold standard FSC value of 0.143; local resolution colouring in three different orientations (left) and of the central section (right) of the reconstructed volume of class 0 presented in the surface representation with the local resolution range scale for **E** non-uniform refinement and **F** local refinement on three regions.

Instead, a 3D heterogenous refinement of the 369,648 particles with maximum alignment resolution set to 10 Å revealed that the sample contained two major populations (Fig. 1B). Class 1, with a population of 108,752 particles, was interpreted as a pseudo helical particle, formed by oligomerization of less than ten β-crustacyanin subunits. Non-uniform 3D refinement of class 1 yielded a map at 3.09 Å resolution, of approximately 175 Å in length and formed by oligomerization of eight β-crustacyanin subunits. Class 0, with a population of 260,896 particles, was also interpreted as a pseudo helical particle but formed by oligomerization of possibly more than ten β-crustacyanin subunits. Non-uniform 3D refinement of class 0 yielded a map at 2.75 Å resolution, with good orientation distribution (Fig. 1C-D).

Local refinement using class 0 particles was performed on three regions achieving local resolution of 2.95 Å, 2.77 Å and 2.96 Å respectively. The three maps thus obtained were combined, using the combine_focused_maps routine of ChimeraX (Pettersen et al., 2021), into a single composite map with local resolution up to 2.5 Å in the central region (Fig. 1E,F).

The α-crustacyanin composite cryo-EM density map (Fig. 2A-D) was fitted with twelve copies of the crystallographic structure of β-crustacyanin reported here (Fig. 2E). The H1 subunits (type I; CRTC) point towards the outer shell of α-crustacyanin, while the H2 subunits (type II; CRTA) are positioned at the inner core of the elongated pseudo helical particle thus obtained (Fig. 2F,G,H).

**Fig. 2.**
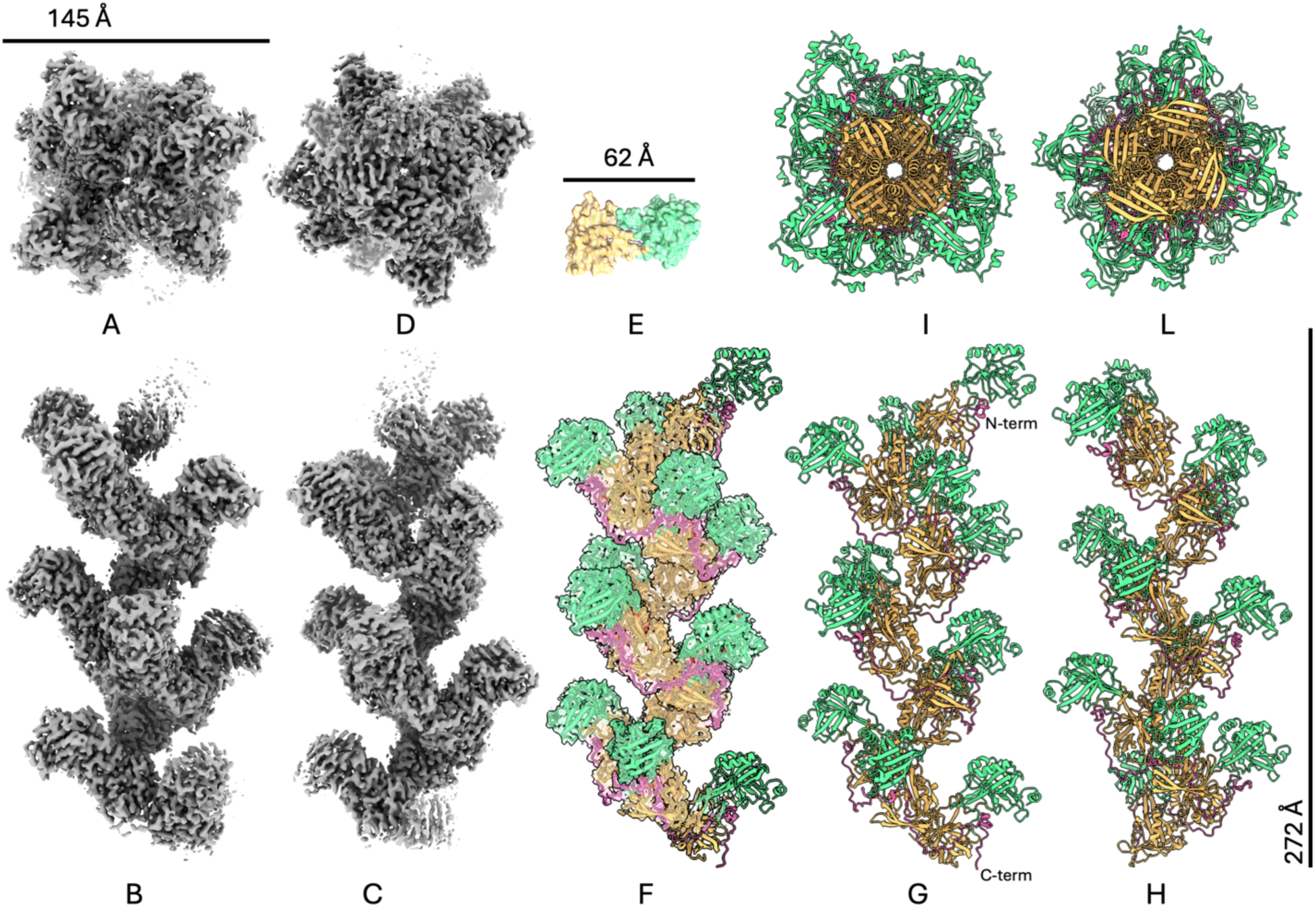
Reconstructed model of α-crustacyanin from *H. americanus* by cryo-EM. representative coulomb density after local refinement for α-crustacyanin at contour level 2.0 σ (**A** top, **B, C** two side views 120 degree apart, **D** bottom). **E** surface representation of β-crustacyanin heterodimer resolved by X-ray diffraction (carton model with colour coding: H1 subunit in green, H2 subunit in beige). **F** fitting of twelve β-crustacyanin heterodimers into the density of α-crustacyanin (HPR chains in pink). **G, H** side views 120 degree apart of the final model. **I,** top view, **L,** bottom view.

In the inner core of α-crustacyanin, each n-1 β-crustacyanin interacts with the n β-crustacyanin via one interaction patch between two independent regions of each H2 subunit. The interaction patch consists of the following residues: Glu145_(n-1)_-Ser14_(n)_, Gln138_(n-1)_-Val15_(n)_, Pro142_(n-1)_-Ser14_(n)_, Gln138_(n-1)_-Asn17_(n)_, Tyr31_(n-1)_-Asp19_(n)_, Ala168_(n-1)_-Phe86_(n)_. Hydrogen bonds are also mediated by a few water molecules (Fig. 4B).

### The heptatricopeptide repeat (HPR) protein

The cryo-EM map revealed additional density indicating the presence of a third, unforeseen, protein component, prompting an extensive analysis on the protein composition of α-crustacyanin sample from *H. americanus*. A Tricine-SDS–PAGE gel (Fig. 3A) confirmed the presence of bands of about 20 kDa corresponding to the two major subunits, H1 and H2 as expected, and additionally few other minor bands. Next, purified α-crustacyanin was denatured in 6 M urea and subjected to size exclusion chromatography under same denaturing conditions (Fig. 3B) followed by Tricine-urea-SDS-PAGE analysis of the collected fractions which confirmed the presence of several high molecular weight polymers between 80 and 120 kDa (Fig. 3C). Mass-spec proteomic analysis of those bands (Fig. 3D) identified the third protein, as the protein Hamer_G005280, labelled as putative tetratricopeptide repeat (TPR) like protein, presenting a repetition of 37 amino acids (heptatrico, HPR) (Fig. 3E). The HPR protein has been modelled on the cryo-EM map as a continuous 12-fold repeat of the 37 conserved amino acids (Fig. 3F,G,H) helical polypeptide wrapping around the α-crustacyanin substructure anticlockwise with the direction N-terminal to C-terminal top to bottom (Fig. 2G,H). The Phe-His dipeptide was used as register mark for correct amino acid sequence assignment (Fig. 3F). Each HPR repeat (1-37) has a defined coil-helix-coil structure. The coil regions comprise of residues 1-10 and 18-37, the helix comprises of amino acids 12-16 of each repeat.

**Fig. 3.**
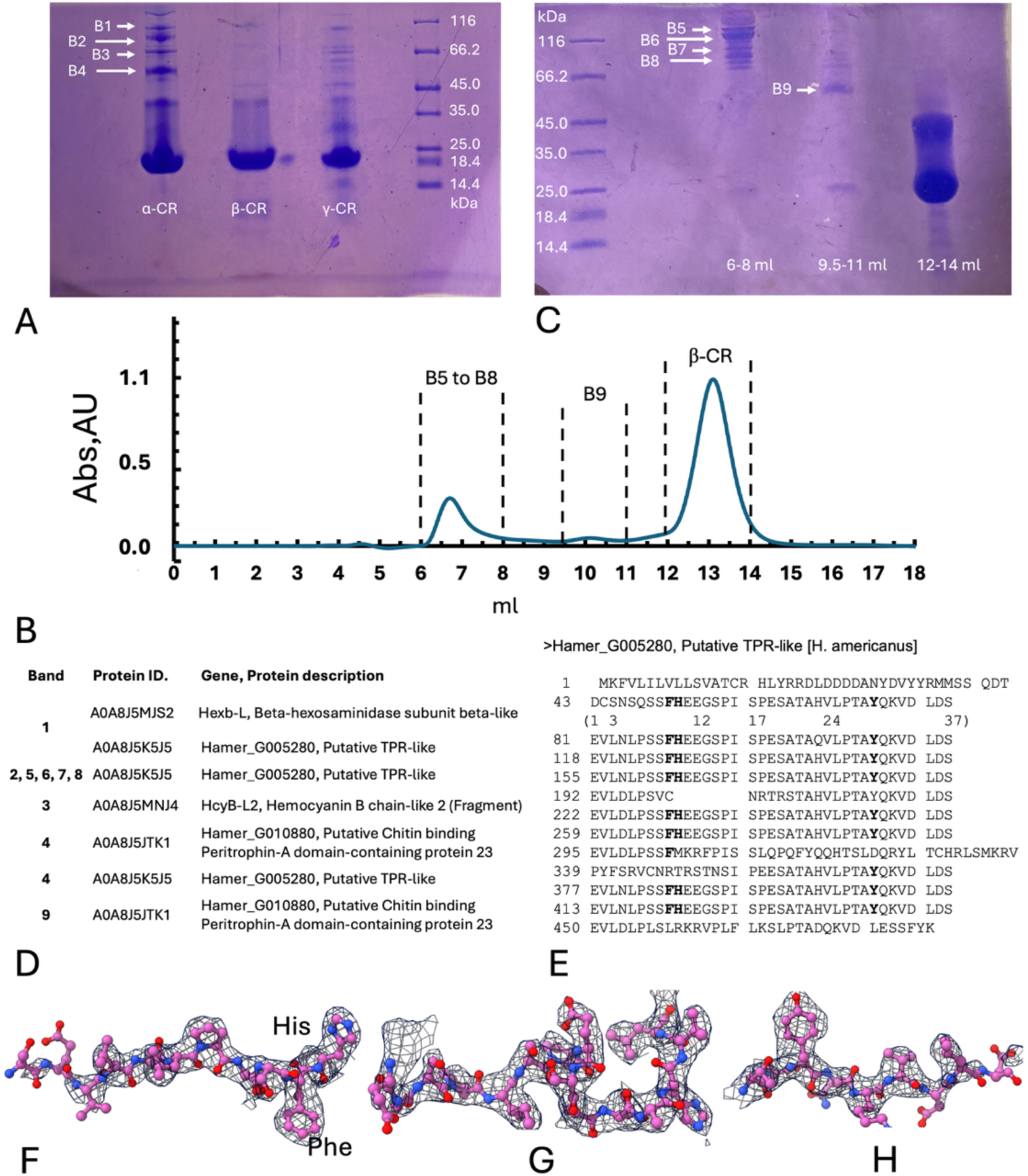
HPR protein characterization. Protein component analysis by SDS-PAGE and proteomic analysis: **A** 10% acrylamide Tricine-SDS-PAGE showing the migration of α-crustacynin (α-CR), β-crustacynin (β-CR) and γ-crustacyanin (γ-CR) samples. It is evident the lack of protein bands B1-4 in the β-crustacyanin and partially in the γ-crustacyanin samples. **B** gel-filtration chromatography in urea denaturing conditions of α-crustacynin sample. **C** 10% acrylamide Tricine-urea-SDS-PAGE of α-crustacynin gel filtration chromatography fractions as indicated in panel **B**. **D** gene assignment of bands 1 to 9 marked in panel **A** and **B**, resulting from *in-gel* tryptic digestion and proteomic analysis. **E** amino acid sequence of the Hamer_G005280 (Uniprot code A0A8J5K5J5), putative TPR-like protein, from *H. americanus*, with the 37 amino acid repeat EVLNLPSS**FH**EEGSPISPESATAHVLPTA**Y**QKVDLDS, with a MW of 3957.32 Da. Key residues Tyr (Y), His (H) and Phe (F) are highlighted in bold; representative densities of the locally refined cryo-EM map contoured at 3.5σ: **F** residues 148-159 **G** residues 159-175 **H** residues 175-184 of the HPR protein.

### The HPR-β-crustacyanin complex unit

Each HPR is bound to one copy of β-crustacyanin heterodimer (Fig. 4A) thus forming a HPR-β-crustacyanin complex unit. The first coil bridges the H2 and the N-terminus region of H1 subunits and insulate the C1’-6’ end ring of AXT1 (Fig. 4C). The helix region, characterized by the SPESATA motif (Ser-Pro-Glu-Ser-Ala-Thr-Ala), holds the N-term of subunit H1 around subunit H2 above the binding site of astaxanthin forming hydrogen bonds between Ser165 and Ser168 of HPR chain and Asp1 of H1. It also interacts with the β-crustacyanin at the interface region between the H1 and H2 subunits above the astaxanthin binding site (Fig. 4D). The second coil region mainly interact with the subunit H2, forming hydrogen bonds and a salt bridge between Arg173 of H2 subunit chain and Asp184 of HPR, leading the chain to the next repeat (Fig. 4E). Supplementary Tables S4, S5 and S6 summarize the main intermolecular interactions between the HPR proteins and the β-crustacyanin subunits.

**Fig. 4.**
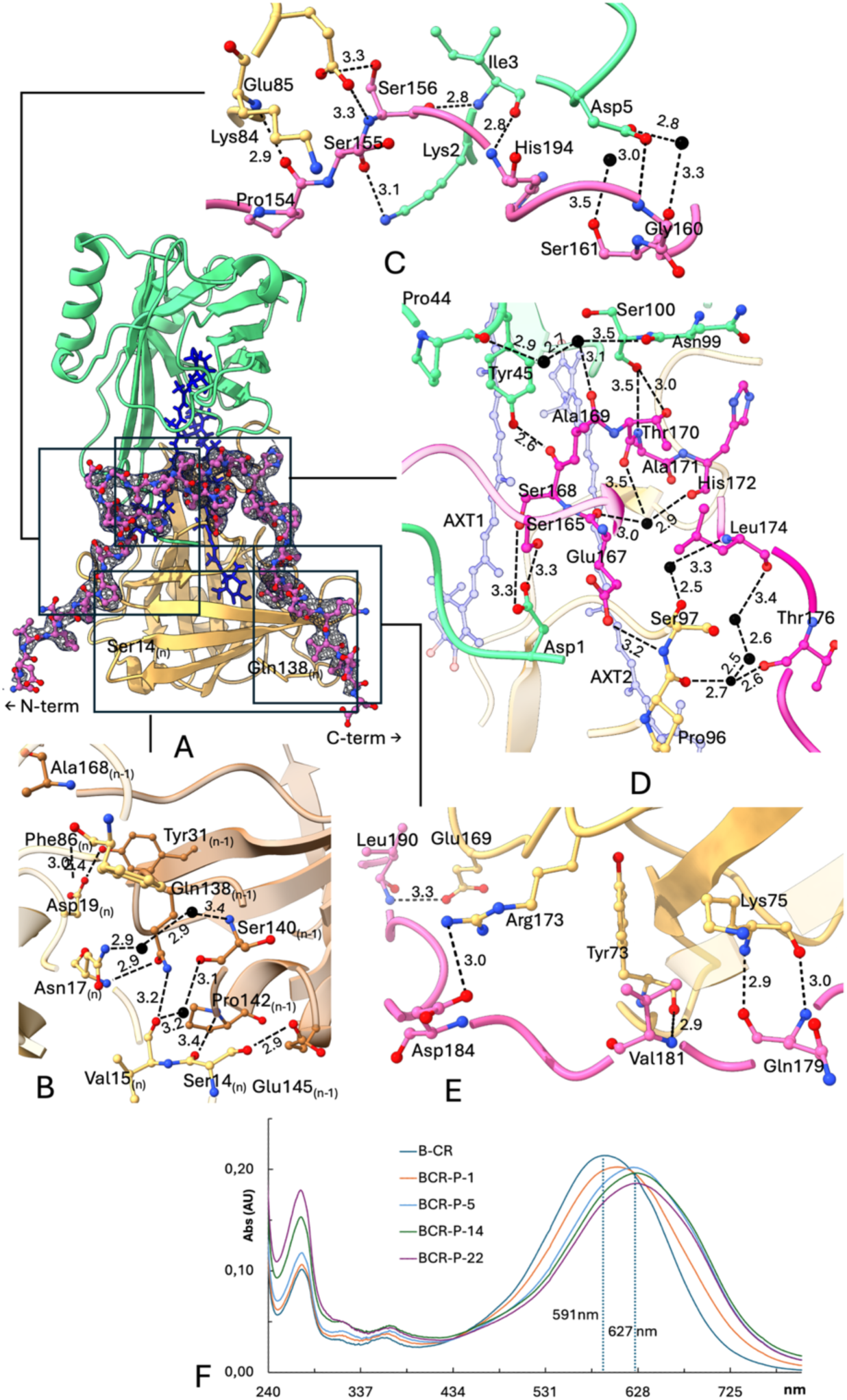
Structural details of the HPR-β-crustacyanin complex unit. **A** reference β-crustacyanin heterodimer (chain H1 green, chain H2 beige, with astaxanthin molecules (blue) and the local refined cryo-EM map of the HPR chain contoured at a 3.5σ level (blue mesh). **B** interaction patch between n (light beige) and n-1 (darker beige) of the H2 subunits. **C, D, E** details of the chemical interactions with the HPR (pink) protein, residues 148-184. Distances are reported in Å. Key residues and waters are depicted with ball-and-stick representation (nitrogen, blue; oxygen, red; waters, black). **F** Absorption spectra recorded during stepwise additions of HPR peptide (P; 4 µL per step, 0.37 mg/mL) to β-crustacyanin (BCR; 0.05 mg/mL) at 6 °C. In the legend *BCR–P–X*, *X* indicates the addition step.

### Spectrophotometric analysis of β-crustacyanin-HPR peptide interaction

To investigate the contribution of the HPR peptide to the bathochromic shift of astaxanthin in β-crustacyanin, in vitro complex formation between the β-crustacyanin unit with a synthetic heptatricopeptide, was followed spectrophotometrically. Stepwise additions of the HPR peptide to freshly purified β-crustacyanin led to a gradual red shift of the absorption spectrum. The *λ_max_* of the astaxanthin progressively increased from 591 nm to 627 nm, indicating successful interaction between the peptide and the protein (Fig. 4F). This assay identifies the contribution of the HPR peptide to the bathochromic shift of astaxanthin in crustacyanins.

### Oligomeric states of α-crustacyanin form

3D classification on single particle cryo-EM data results into two distinct classes based on particle length, namely class 0, describing a long oligomeric complex, and class 1 describing a smaller complex (Fig. 1B). Further 3D classification (Fig. 5A) on each of these two populations using heterogenous refinement reveals the oligomeric composition of α-crustacyanin to be between eight to ten HPR-β-crustacyanin complex units for class 1 (Fig. 5B) and from eleven to sixteen for class 0 (Fig 5C).

**Fig. 5.**
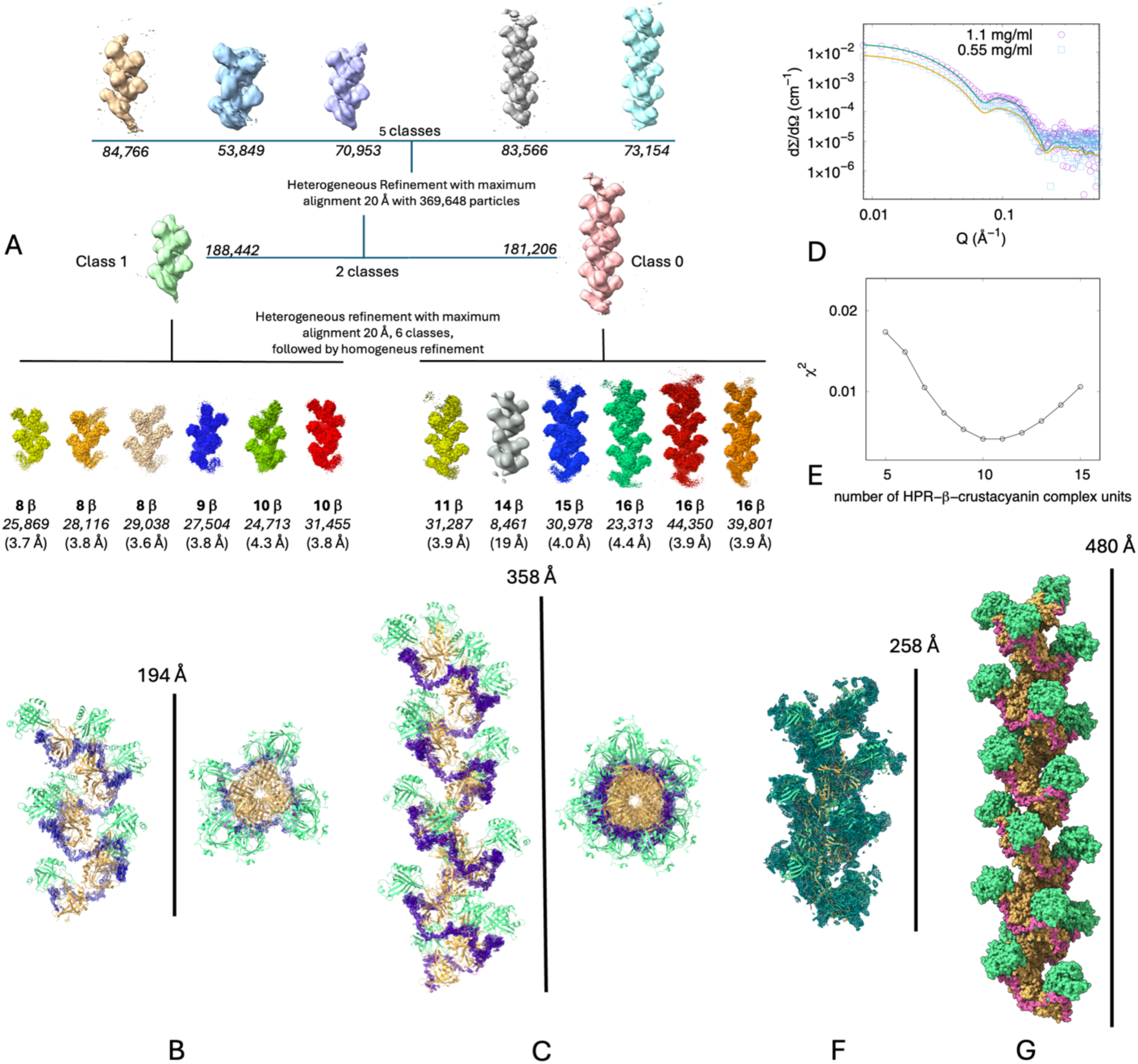
Oligomeric model of α-crustacyanin from *H. americanus* evaluated by cryo-EM, small angle X-ray scattering and X-ray crystallography. **A** heterogenous refinement with maximum alignment at 20 Å, with five (top) and two classes (bottom), then further divided in six classes. Number of HPR-β-crustacyanin complex units is indicated in bold, number of particles in italic and achieved resolution in brackets. **B** representative structure of class 1 (8-mer) fitting the map post heterogeneus refinement (resolution 3.09 Å). **C** representative structure of class 0 (16-mer) fitting the composite map (resolution 3.9 Å). **D** profile-fitting results of SAXS data at two different concentrations using 10-mer and 11-mer α-crustacyanin models. **E** χ^2^ model fitting values as function of the number of HPR-β-crustacyanin complex units. **F** composite omit Fourier difference electron density map (lavender) contoured at 1 sigma overlapping the refined crystallographic structure resolved at 6.32 Å resolution using the cryo-EM structure as model. **G** extended molecule of α-crustacyanin by applying the crystallographic symmetry operation (−1,+1,0) to obtain an elongated filament of α-crustacyanin complexes.

The SAXS curves (Fig. 5D, E) could be fitted with single input models of α-crustacyanin of different length. The lowest χ^2^ values correspond to theoretical fitting curves with models of oligomeric number from ten to twelve of HPR-β-crustacyanin complex units (Fig. 5E) thus confirming the pseudo helical architecture matching the cryo-EM structure.

Similarly, the crystallographic structure solution was attempted at 6.32 Å resolution with molecular replacement using as inputs the cryo-EM model of α-crustacyanin with an oligomeric length from four to twelve HPR-β-crustacyanin complex units. The best solution with an α-crustacyanin model of eleven units had translation- function Z-scores of 17.3 for the fitted molecule, an overall log-likelihood gain of 241.8 (Table S3). Other possible solutions were discarded upon direct inspection of crystal lattice defects using COOT (Emsley et al., 2010). The calculated Matthews coefficient was of 4.5 corresponding to 72.9 % of solvent content, similar to the one of β-crustacyanin heterodimer (Table S1). The structure was refined using REFMAC Servalcat (Yamashita et al., 2021) within the CCPEM suite (Wood et al., 2015) and PHENIX refine (Adams et al., 2010) resulting in a final R_work_ and R_free_ of 27.1% and 37.3%. The final average NCS CC at full resolution with 11 operators calculated using the PHENIX find_ncs protocol (Terwilliger, 2002) was only 0.5, thus confirming the pseudo helical nature of α-crustacyanin architecture (Fig. 5F). Expanding the α-crustacyanin crystallographic structure beyond the asymmetric unit by applying the crystal symmetry operations, the propensity of α-crustacyanin to form oligomers of oligomers is clearly revealed (Fig 5G). The same inter-oligomeric interactions could potentially happen between any combination of class 0 and class 1 particles thus forming complexes with repeated units beyond eleven. Gel filtration of the α-crustacyanin complex resulted in an approximate average molecular weight of around 762 kDa (Cedri et al., 2026). Given the calculated molecular weight of the HPR-β-crustacyanin unit of 47.5 kDa, we could estimate a molecular complex of sixteen HPR-β-crustacyanin units.

Considering that elongated shaped particles tend to elute faster in SEC, the approximate average molecular weight could be smaller, however. Taken together, the crystallographic, SAXS and cryo-EM data confirm that the α-crustacyanin complex presents a pseudo helical architecture of α-crustacyanin complex in crystallo and in solution, of variable oligomeric length from eight to sixteenth HPR-β-crustacyanin units.

### Binding of astaxanthins within the α-crustacyanin

In the α-crustacyanin form the binding astaxanthin sites result from the interactions between the two lipocalin subunits H1 and H2 and with the HPR protein. Each HPR-β-crustacyanin heterodimer binds two astaxanthin molecules, AXT1 and AXT2, in a *6-s-trans* conformation with the end rings coplanar with their polyene chains, which remain 6.7 Å apart (Fig. 6, centre-top). In each astaxanthin molecule, the C1-6 ring is in the lipocalin calyx, while the C1’-6’ ring sits at the interface of subunits H1 and H2. AXT1 has its C1-6 ring bound to H1 calyx, and AXT2 to H2 calyx (see Fig. 6). The repeated α-helix region of the HPR protein interacts with the H1 and H2 subunits at the centre of the complex, above the astaxanthin binding sites at the surface-exposed hydrophobic patch (Fig. 6, centre-middle). The end ring C1-6 of AXT1, is within hydrogen bond distance from the side chain of residues of the H1 subunit Gln46 and Asn54, and constrained within Ile95, Phe134, and Phe136 (Fig. 6A). The end ring C1’-6’ is in proximity of Phe86, His90, Pro102, Phe126 and Tyr128. The C1’-6’ end ring carbonylic oxygens are found at 3.2 Å from the N atom of His90 imidazole groups (Fig. 6C).

**Fig. 6.**
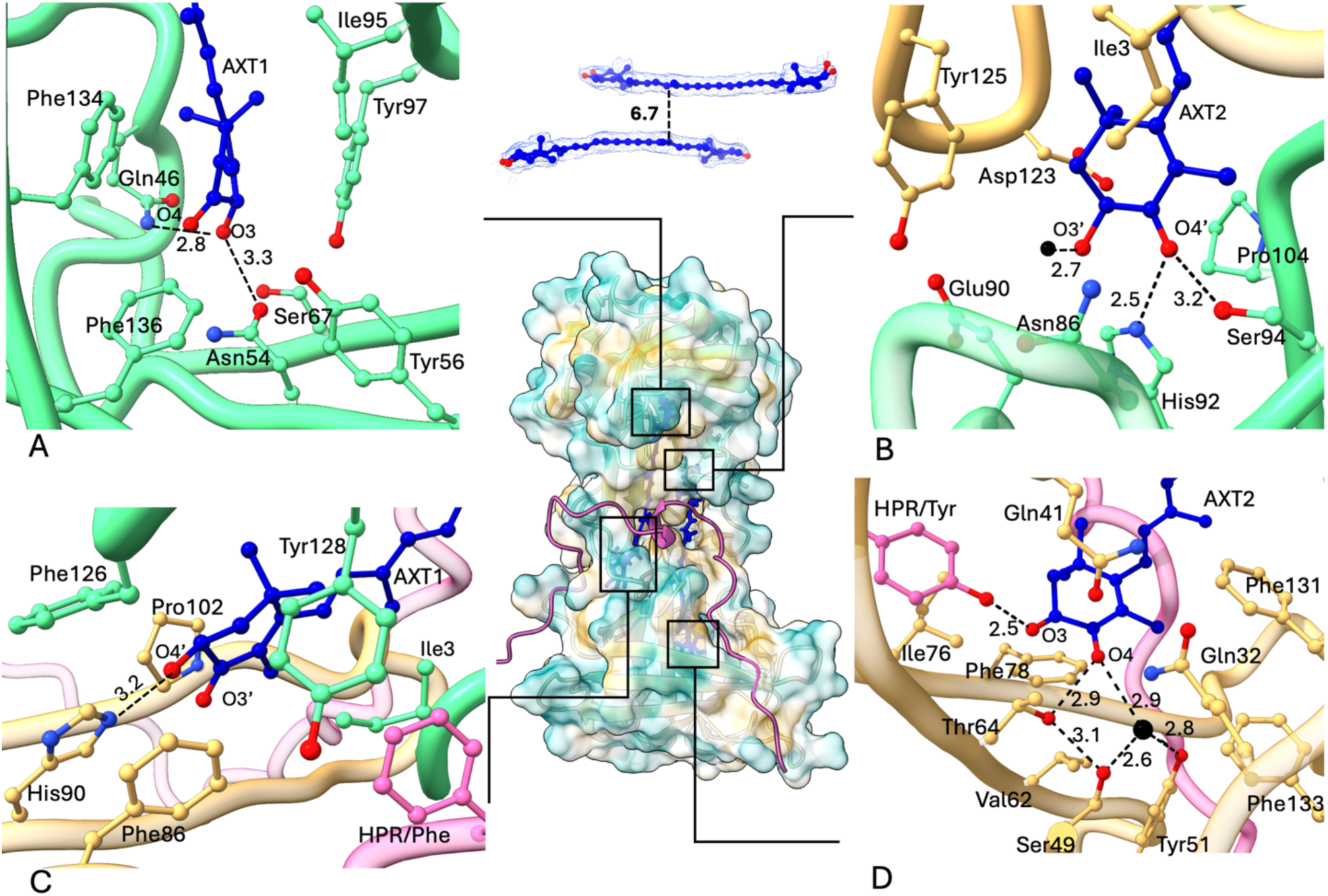
Astaxanthin (AXT) binding sites in α-crustacyanin from *H. americanus* and spectrophotometric analysis of the β-crustacyanin–HPR peptide interaction as a function of peptide concentration and temperature. **centre, top,** representative two bound AXT molecules depicted in blue colour in ball-and-stick format with Coulomb density at 3.5σ level. **centre, middle** hydrophobic surface representation of the HPR-β-crustacyanin with the HPR protein in pink ribbon. Colour coding for the hydrophobic surface is from dark cyan for most hydrophilic through white to dark goldenrod for most hydrophobic. The subunit calyxes are depicted: H1 in green and the H2 in beige. The averaged distances in Å for the binding interactions are reported in: **A** ring C1-6 AXT1 binding site; **B** ring C1’-6’ AXT2 binding site; **C** ring C1’-6’ AXT1 binding site; **D** ring C1-6 AXT2 binding site.

Conversely, for the AXT2 molecule, the end ring C1-6 interacts with the main chain of H2 subunit residues Thr64, Phe131, Tyr51, and Ile76 (Fig. 6D). Additionally, the repeated coil region of the HPR protein is positioned such to bring the phenolic ring of the HPR/Tyr at close distance of 2.5 Å from the hydroxy-group of the end ring C1-6 of the astaxanthin molecule (Fig. 6D). The end ring C1’-6’ is in proximity of His92, Pro104, and Tyr125. The C1’-6’ end ring carbonylic oxygen is found at 2.5 Å from the N3 atom of His92 imidazole group (Fig. 6B).

When superposing the astaxanthin binding sites of the β-crustacyanin crystallographic form and the α-crustacyanin cryo-EM structure, the water molecules observed near the C1-6 and C1’-6’ of the AXT2 molecule appear to be conserved (Fig. S3A,B,C).

One water molecule is found within hydrogen bond distance from Tyr51(H2) and Ser49(H2) and the O4 from the carbonylic group at the end ring C1-6 of the AXT2 (Fig. S3A,C). The second water molecule forms hydrogen bond exclusively with the O3’ from the carboxylic group of the C1’-6’ ring of AXT2 (Fig. S3B).

Water molecules present in the proximity of the astaxanthin binding sites would enable charge transfer from the amino acid side chains to the keto and the hydroxy group of the astaxanthin end rings, thus promoting the polarization of the carotenoid (Britton et al., 1997).

The Ala-Thr-Ala region of the HPR helix is directly above the methyl groups of central polyene regions of two astaxanthin molecules (Fig. 6, centre), holding the phenyl rings of Phe6(H1), Phe101(H1) and Phe99(H2) in place between the polyene carbon chains, thus insulating the polyene chain from the exocuticle carbonate structure (Fig. 7E). The complete polyene chain insulation and the presence of the HPR/Tyr phenolic ring next to the hydroxy-group of the C1-6 end ring of the astaxanthin molecule, bound within the H2 calyx (Fig. 6D), are the only two significant differences observed in the binding of astaxanthin between α- and isolated β- forms.

**Fig. 7.**
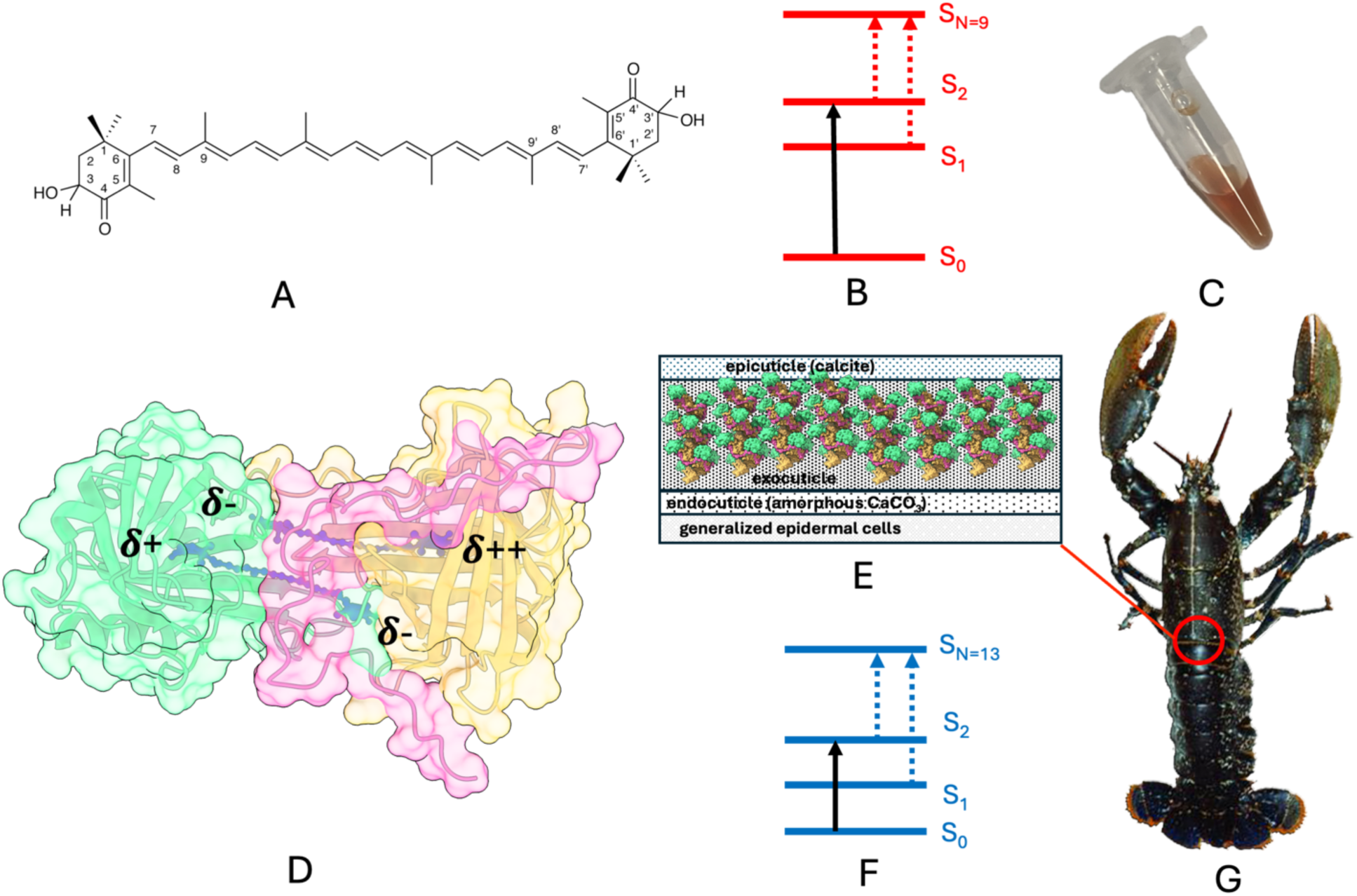
Chemical basis of a lobster exocuticle blue colouration. **A** chemical structure of the chromophore astaxanthin with atom numbering. **B** schematic representation of the four states energy-level model (S_0_, S_1_, S_2_ and S_N_) of a carotenoid molecule with N=9 (unbound form, S_0_ → S_2_ wide gap). **C** typical red colour of an ethanol solution of astaxanthin. **D** HPR-β-crustacyanin units of α-crustacyanin complex containing astaxanthin with polarization of the end rings indicated. **E** model of a lobster cuticle with positioning of α-crustacyanin complex in the exocuticle. **F** schematic representation of the four states energy-level model (S_0_, S_1_, S_2_ and S_N_) of a carotenoid molecule with N=13 (bound form, S_0_ → S_2_ narrow gap). **G** representative dark-blue uncooked lobster carapace.

The binding of astaxanthin molecules is consistent in all the twelve HPR-β-crustacyanin repeat complex units observed within the α-crustacyanin complex. In fact, the distances between the end rings and the key residues within the calyxes of the H1 and H2 subunits are the same within the experimental error (Table S7). The geometry of the astaxanthin molecules is preserved within the β-crustacyanin crystal structure and the α-crustacyanin cryo-EM structure without any evidence of major distortions. In fact, the r.m.s. deviations are in the range 0.3 Å to 1 Å (Tables S8 and S9) showing consistency in the binding mode of astaxanthin both in crystallo and in solution.

It is worth mentioning that in the natively purified EPD-related blue carotenoprotein-1 the two carotenoids are specifically bound to the heterodimer interface, where the polyene chains are aligned in parallel to each other at 6.7 Å distance like in the HPR-β-crustacyanin complex, although the two complexes are evolutionary and structurally unrelated (Kawasaki et al., 2023).

### Quantum chemical calculations

Quantum chemical calculations were performed to estimate the contribution of exciton coupling to the bathochromic shift. To this end, the most accurate approach is to consider the interactions between the two carotenoids in each AXT dimer explicitly, as demonstrated in previous work (Strambi & Durbeej, 2009). This “supermolecule” approach is based on the fact that any monomeric excited state Ψ^∗^ splits into two excited states 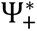 and 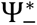 as result of exciton coupling, each with a distinct energy and transition dipole moment. Accordingly, the contribution of exciton coupling to the bathochromic shift can be calculated as half the energy difference between these two states 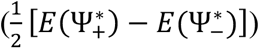. All calculations were carried out using density functional theory methods as implemented in the Gaussian 16 suite of programs (Frisch et al., 2009). First, the geometries of each of the 12 AXT dimers in the cryo-EM structure were optimized with the range-separated, dispersion-including ωB97X-D functional (Chai & Head-Gordon, 2008) in combination with the 6-31G(d,p) basis set. In order to obtain physically relevant geometries, the optimizations made use of a few constraints on the interatomic distances and the intramonomeric dihedral angles, as shown below. For each dimer, the specific values of the constraints were kept at those shown by the cryo-EM structure.

| Constrained interatomic distances |  |  |
| --- | --- | --- |
| C13(A)–C13'(B) | C12(A)–C12(B) | C12'(A)–C12'(B) |
| C6(A)–C6(B) | C13(A)–C13(B) | C11'(A)–C11'(B) |
| C7(A)–C7(B) | C14(A)–C14(B) | C10'(A)–C10'(B) |
| C8(A)–C8(B) | C15(A)–C15(B) | C9'(A)–C9'(B) |
| C9(A)–C9(B) | C15'(A)–C15'(B) | C8'(A)–C8'(B) |
| C10(A)–C10(B) | C14'(A)–C14'(B) | C7'(A)–C7'(B) |
| C11(A)–C11(B) | C13'(A)–C13'(B) | C6'(A)–C6'(B) |
| Constrained intramonomeric dihedral angles |  |  |
| C5(A)–C6(A)–C7(A)–C8(A) | C5'(A)–C6'(A)–C7'(A)–C8'(A) | C5(B)–C6(B)–C7(B)–C8(B) |
| C5'(B)–C6'(B)–C7'(B)–C8'(B) |  |  |

For each of the resulting dimer geometries, the energies of the two relevant excited states needed to quantify the exciton coupling by the supermolecule approach were obtained by performing single-point calculations with the ωB97X-D, CAM-B3LYP (Yanai et al., 2004) and B3LYP density functionals, again with the 6-31G(d,p) basis set. The corresponding results, shown in Fig. S4, reveal that the contribution of exciton coupling to the bathochromic shift is consistently amounting to no more than 0.04-0.05 eV for each AXT dimer. Here, it should be emphasized that these calculations were performed without accounting for potential effects from the protein. However, based on the observation that the coupling previously calculated in the same way for just a single AXT dimer (0.04 eV (Strambi & Durbeej, 2009)) agrees very well with the experimental value of 0.03 eV obtained from measurements on *H. americanus* β-crustacyanin, (Christensson et al., 2013) this appears to be a reasonable approximation.

## Discussion

### The molecular basis of the bathochromic shift from 591 nm to 631nm

The unbound dilute astaxanthin (Fig. 7A, C) has a λ_max_ = 472 nm in hexane, while it has a λ_max_ = 591nm when bound in the β-crustacyanin form and 631nm when bound in the α-crustacyanin complex. Comparison of the structures of both *H. americanus* β-crustacyanin and α-crustacyanin allows to pinpoint at atomic level the contributors responsible for the bathochromic shift from 591 nm to 631nm.

The determinant of the colour of a specific carotenoid is the total count (N) of conjugated C=C plus C=O bonds within the molecule (Fig. 7A) (Polívka & Sundström, 2004). The spectroscopic characteristics of neutral carotenoids can be described by using a three-state model comprising the ground state, S0, and two excited singlet states labelled S1 and S2. Selection rules prohibit the S0 → S1 transition, while the transition from S0 to the second excited state, S2, is strongly permitted (Fig. 7B). The energy associated with the S0 to S2 transition decreases when N increases, causing a corresponding shift in the absorption band towards longer wavelengths. The corresponding complementary visible colours can be associated using the Munsell’s colour wheel (Ferrari et al., 2012; Munsell, 1912). For neutral carotenoids with very long conjugated π-electron chain lengths, the spectral onset of the S0 to S2 transition falls within the blue-green region of the visible spectrum, contributing to the characteristic yellow, orange, and red hues in organic solvents (Polívka & Sundström, 2004). Astaxanthin follows these rules. In hexane, free astaxanthin has a relaxed geometry, with the end ring out of the plane of the polyene chain, resulting in N=9, as proved by chemical crystallography studies on astaxanthin and its derivatives (Bartalucci et al., 2007, 2009; J. R. Helliwell, 2010; M. Helliwell, 2008; M. Helliwell & Helliwell, 2007; Polívka et al., 2010). The binding of astaxanthin to the β-crustacyanin scaffold contributes to the bathochromic shift to 591 nm observed here, resulting in three key perturbations to the orbital state of astaxanthin. The two carotenoid binding sites force the marked bowing of each AXT molecule (Fig. 6, centre, top), with the end rings of astaxanthin held coplanar in an S-trans orientation with the polyene chain (Fig. 6), thus extending the polyene conjugation to N=13. Quantum calculations have quantified that these geometrical regularisations of the astaxanthin geometry contributes to increase the bathochromic shift by 20-30 nm (Durbeej & Eriksson, 2004; Gamiz-Hernandez et al., 2015). The end-ring keto-oxygens are positioned in an asymmetric environment within hydrogen bonding distance from polar residues (Fig. 6) and from four bound waters observed in the β-crustacyanin structure of *H. americanus* reported here. Protonation of the conjugated carbonyl groups, hydrogen bonding to the histidine residue and water, also contribute to the bathochromic shift by about 100 nm, as indicated by mechanical calculations (Begum et al., 2015; Durbeej & Eriksson, 2003, 2006, 2004; Gamiz-Hernandez et al., 2015; Strambi & Durbeej, 2009). The protonation mechanism is also supported by in vitro recombination studies of recombinant H1 and H2 apo subunits with astaxanthin, obtaining in both cases an absorption peak at 580 nm in *H. gammarus*, when only one astaxanthin molecule would bind to a single apo subunit within the calyx, with minimal difference in the absorption peak of the reconstituted β-crustacyanin (Ferrari et al., 2012).

Exciton coupling, or the coupling of the excited states of two identical chromophores in proximity (Berova et al., 1999), has been used to explain the large bathochromic shift (Ilagan et al., 2005; Neugebauer et al., 2011; van Wijk et al., 2005) but was lately discarded (Christensson et al., 2013; Gamiz-Hernandez et al., 2015; Strambi & Durbeej, 2009). The β-crustacyanin crystallographic structure reported here by itself does not provide any additional evidence to support a contribution coming from exciton coupling. This is, even though the side chains of Phe6 (H1), Phe101(H1) and Phe99(H2) are now found between the polyene chains. Such arrangement could possibly increase the coupling of the conjugated system of the carotenoids through these aromatic side chains, but instead no additional bathochromic effect is observed. In other words, β-crustacyanins from *H. gammarus* and from *H. americanus* have a reported an absorption peak in the 585 – 591 nm range with or without aromatic side chains placed between the astaxanthin molecules.

In vitro complex formation between the β-crustacyanin unit with a synthetic heptatricopeptide to form the β-crustacyanin-HPR unit (Fig. 4F), confirms the HPR protein, in concert with crustacyanins, as major contributor in tuning carapace colours up to 627 nm thus inducing further bathochromic shift.

There are several contributions of the HPR protein to the bathochromic shift of 40 nm from 591nm in isolated β-crustacyanins to 631 nm in α-crustacyanins when forming the β-crustacyanin-HPR complex repeat (Fig. 6, 7) within the α-crustacyanins architecture (Fig. 5). The HPR protein increases the polarization of the end-ring keto-oxygens. In fact, a phenolic ring of the HPR/Tyr is found within hydrogen bond distance (2.5 Å) from the hydroxy group at the C1-6 end ring of the AXT2, bound within the calyx of the H2 monomer where Tyr51 side chain is also present (Fig. 6D). Both Tyr residues increase the positive partial charge via a network of hydrogen bonds to a single end of one astaxanthin due to the weak acidic nature of a phenolic group which is likely to be protonated at pH 7. The benzene ring of the HPR/Phe is within 4 Å from the hydroxy group of the other end ring of the astaxanthin molecule bound in the H2 monomer (Fig. 6C and Fig. S3D) thereby insulating this site from the outside environment. In these conditions the His residues in proximity of the C1’-6’ for the astaxanthin end ring (Fig. 6B,C) would induce the enolate formation as proposed by Begum et al. (Begum et al., 2015). In forming the β-crustacyanin-HPR complex, the HPR helix region completely insulates the polyene chains of two astaxanthin molecules thus avoiding interference from the embedding environment (Fig. 7E). All these contributions would favour astaxanthin polarization (Fig. 7D) decreasing the energy of the C=O antibonding orbitals that can thus participate more efficiently in the S0 stability of the astaxanthin molecule in α-crustacyanin. This would lead to a significant decrease of the energy gap between the ground S0 and first excited electronic state S2 (Fig. 7F). Furthermore, the HPR/Tyr residue is positioned next to a single end ring - the one of the AXT2 - out of the four end rings of AXT1 and AXT2. The fact that the complete spectra of both astaxanthins is red shifted to 631 nm with unimodal distribution, and no residual peak or shoulder is observed at 591 nm (Fig. 4F), meaning that the two astaxanthin molecules are equally affected. This implies that the additional positive partial charge induced by the HPR/Tyr to AXT2 is also perceived by AXT1 molecule. In fact, the two astaxanthins within the β-crustacyanin-HPR complex repeat motif experience a slightly different ligand environment, while generating a single absorption peak in α-crustacyanins, so de facto behaving as a single chromophore. The exciton coupling mechanism was therefore re-examined. We performed quantum chemical calculations on each of the 12 astaxanthin dimers present in the cryo-EM structure. Encouragingly, these results confirm that the exciton coupling, generated by the interaction of the π-electron systems, accounts for no more than 0.04-0.05 eV for each astaxanthin dimer (Fig. S4) i.e. about 10% of the entire bathochromic shift from 480 nm (2.583 eV) to 591 nm (2.098 eV). This means that the contribution of exciton coupling to the bathochromic shift is not much larger for the α- than for the β-crustacyanin. It remains to be investigated how the two astaxanthins could possibly achieve a unimodal distribution. The limitation in the interpretation of our results is that, as it appears from the architecture of α-crustacyanin reported here, there are several possible contributors to the bathochromic shift. This calls for a detailed investigation on the role of molecular dynamics and /or quantum chemical calculations as attempted by Loco et al. (Loco et al., 2018) with the β-crustacyanin crystal structure from *H. gammarus* (Cianci et al., 2002) to quantify these contributions. Broadband transient absorption spectroscopy could be attempted to investigate the singlet fission process in crustacyanin bound astaxanthin as suggested by Musser et al. (Musser et al., 2015).

### The HPR-crustacyanin protein complexes in other marine invertebrates

Our studies demonstrate that α-crustacyanin complex, ex vivo purified from the American lobster shell, is a complex with a variable number of copies of the astaxanthin binding β-crustacyanin and the heptatricopeptide repeat (HPR) protein. Here, as a constituent of α-crustacyanins from *H. americanus*, the HPR protein adds other level of complexity. It promotes the ordered organization of multiple copies of the β-crustacyanin-HPR complex repeat motifs, resulti in a darker and more intense blue hue of the lobster carapace (Fig. 7G).

Repeat proteins have been previously reported in many organisms. The tetratrico (TPR) family is widespread in eukaryotes including *P. falciparum*, *T. gondii* and *H. sapiens* and primarily involved in RNA metabolism, with possible function similar to that of octatrico peptide repeat (OPR) and pentatrico repeat proteins (PPR) proteins (Hillebrand et al., 2018). In *Arabidopsis* mithocondria PPR play an important role in RNA processing and transcript stabilization (Colcombet et al., 2013). The OPR family, is more prevalent in single-celled photosynthetic algae, *e.g. Chlamydomonas reinhardtii* where it is involved in the translation initiation of the PsaB subunit of photosystem I (Rahire et al., 2012).

A BLAST search (Altschul et al., 1997) using the HPR sequence (XP_042225416.1), reveals the presence of the SPESATA motif protein also in other aquatic species like the Australian red claw crayfish or *Cherax quadricarinatus* (XP_053656413.1), and the red swamp crayfish *Procombarus clarkii* (XP_045620142.1). The identification of this protein, together with the reported presence of crustacyanin proteins in *Cherax quadricarinatus* (Luquet et al., 2009) and *Procombarus clarkii* (Chen et al., 2021; Milicua et al., 1986), would suggest a similar architecture for α-crustacyanin in these species.

## Conclusions

Shell colouration deployment and adaptation are essential for the survival and replication of living organisms. Chemical and biological understanding of this process starts from a detailed description of the interactions between chromophores and binding proteins followed by quantum chemical calculations. For these reasons it is essential to have a detailed description of all the possible interactions that occur at the atomic level. In this study such level of detail was achieved by harnessing data from macromolecular crystallography, cryo-EM reconstruction and SAXS curve fitting. Astaxanthin complexes could cover the entire visible colour palette as natural colourants thus offering a versatile alternative to synthetic dyes for the food industry (Landim Neves et al., 2021). In addition, the structural and spectroscopic properties of astaxanthin or similar carotenoids also hold potential interest for their applications as light-absorbing pigments in artificial systems aiming to broaden the spectrum of solar energy absorption (Hashimoto et al., 2015; Polívka & Frank, 2010). Finally, our findings provide a blueprint for designing nutraceutical food pigments and bio-inspired light-harvesting mimetics with tuned light absorption.

## Materials and Methods

α- and β-crustacyanin proteins were purified ex vivo from frozen tails of *H. americanus* by as previously reported and assessed for mono dispersity and stability prior to crystallization, cryo-EM and SAXS experiments (Cedri et al., 2026). Diffraction data were collected at the European Synchrotron Radiation Facility (ESRF, Grenoble, France) at beamline ID30B (McCarthy et al., 2018). Cryo-EM experiments were performed at the Imaging Facility of the City University of New York Advanced Science Research Center (New York, US). Small Angle X-ray scattering (SAXS) experiments were performed at the Elettra Synchrotron (Trieste, Italy), at the Austrian SAXS beamline (Amenitsch et al., 1995). Structural data were analysed using the cryoSPARC (Punjani et al., 2017), PHENIX (Adams et al., 2010), CCP4 (Winn et al., 2011), GENFIT (Spinozzi et al., 2014) and CRYSOL (Franke et al., 2017) software packages. Quantum chemical calculations were carried out using density functional theory methods as implemented in the Gaussian 16 suite (Frisch et al., 2009).

### α-crustacyanin structure determination and validation

The structure was solved by molecular replacement using PHASER (McCoy et al., 2007) after extensive search using (5-14)-mers models derived from the cryo-EM solution of α-crustacyanin. The top TFZ scoring solutions were cross-checked for packing errors using COOT (Emsley et al., 2010). The low-resolution structure was refined using LORESTR (Nicholls et al., 2017) within the CCP4 suite (Winn et al., 2011). Data refinement statistics are summarized in Table S1.

### β-crustacyanin structure determination and validation

The structure was solved by Molecular Replacement using PHASER (McCoy et al., 2007) with one monomer of the native form as a starting model of *H. gammarus* β-crustacyanin (PDB: 1GKA (Cianci et al., 2002). The manual fitting of the side chains and solvent molecules into electron density maps were performed using COOT(Emsley et al., 2010) and the PHENIX suite (Adams et al., 2010), while monitoring R_work_, R_free_, Ramachandran plot (Ramachandran et al., 1963) and related geometrical parameters using Molprobity (C. J. Williams et al., 2018) within the PHENIX suite. The model was finally refined with REFMAC5 (Murshudov et al., 2011). Data refinement statistics calculated using the crystallographic validation tools within PHENIX (Urzhumtseva et al., 2009)are summarized in Table S1.

### Gel filtration and protein component analysis of the α-crustacyanin

Gel Filtration Chromatography was performed on Superose ™ 12 10/300 GL column pre-equilibrated with 6 M urea 50 mM potassium phosphate buffer pH 7.0. α-crustacyanin (8 mg/mL), previously denatured in the same buffer, was loaded onto the column at a flow rate of 0.4 mL/min (Fig 3B). Fractions of the α-crustacyanin sample corresponding to distinct elution peaks were collected and analysed in Tricine-urea-SDS-PAGE (Fig. 3C). The bands were subjected to *in-gel* tryptic digestion and mass-spec proteomic analysis. Protein identification was performed against Uniprot database of *H. americanus* (UP000747542, ID6706, 24351entries, date created: 17052023, date downloaded: 15082023). The analysis was performed as a service by the Proteomics Core Facility at European Molecular Biology Laboratory, (EMBL) Heidelberg, Meyerhofstr. 1-69117 Heidelberg, Germany.

### Cryo-EM grid preparation and cryo-EM data collection

Cryo-EM experiments were performed at the Imaging Facility of the City University of New York Advanced Science Research Center (New York, US). Grids were glow-discharged using the Fischione M1070 NanoClean to make their surface hydrophilic.

For cryo-EM sample preparation ∼4 μL of sample were deposited on grids (Quantifoil R1.2/1.3 Mesh 300, Germany) in 100% humidity and 10 °C and, which were then blotted with filter paper for 2 s using the Vitrobot Mark IV (ThermoFischer™) and quickly plunge frozen in liquid ethane to allow the formation of a thin layer of amorphous ice with the particles embedded within. 10,114 movies (30 frames each) were recorded using a ThermoFischer Titan Halo 80-300 TEM™ microscope equipped with X-FEG high-brightness gun at 300 kV and a Gatan™ K3 Summit Direct Detection Camera operated in electron counting mode, via the SerialEM v. 3.8 control software (Mastronarde, 2005). Nominal magnification was set to 37k for a calibrated pixel size of 0.8465 Å. Total dose of 85.70 e^−^/Å^2^ was spread over four seconds of exposure time resulting in a calibrated dose of 1.07 e^−^/Å^2^/frame. Defocus values were set in a range between -0.8 μm and -2.2 μm, with 0.2 μm steps. Data were collected at a flat stage tilt-angle.

### Image processing

All micrographs were inspected and motion-corrected using WARP (Tegunov & Cramer, 2019). After CTF estimation and correction, particles were picked using an automated protocol embedded in WARP with a box size of 800 pixel. Picked particles were imported to cryoSPARC v.3.2. (Punjani et al., 2017), installed on a SingleParticle™ GPU workstation equipped with four NVIDIA RTX 3090 Graphics Cards and 512 GB ECC Registered DDR4 DRAM (Department of Agricultural, Food and Environmental Sciences, Università Politecnica delle Marche, Ancona, Italy). The final refinement protocol consisted of a non-uniform 3D refinement of 260,896 particles yielding a map at the resolution of 2.75 Å resolution. Local resolution refinement performed on three separate regions achieved local resolutions up to 2.58 Å (see Results section). Global resolution and focused resolution were assigned in cryoSPARC according to the gold-standard FSC (Pintilie & Chiu, 2021).

Heterogenous refinement at 20 Å maximum alignment resolutions were attempted to delineate filament subunit composition, followed by homogeneous refinements (Fig. 5).

### Cryo-EM model reconstruction and validation

The density maps were resized and recentred using the map_box routines within PHENIX suite (Adams et al., 2010). Multiple copies of the crystallographic structural model of *H. americanus* β-crustacyanin, previously solved, were docked into cryo-EM composite local density maps using ChimeraX (Pettersen et al., 2021). Then iterative rounds of manual refinement in COOT v.0.9.5 (Emsley et al., 2010), real_space_refine routine (Afonine et al., 2018a), and finally with REFMAC Servalcat (Yamashita et al., 2021) within the CCPEM suite (Yamashita et al., 2021) followed. Model validation was performed using MolProbity (C. J. Williams et al., 2018) available within PHENIX suite (Afonine et al., 2018b). Data collection and refinement statistics are reported in Table S2. Figures were prepared using ChimeraX (Pettersen et al., 2021).

### Small-angle X-ray scattering

Small Angle X-ray scattering (SAXS) experiments were performed at the Elettra Synchrotron (Trieste, Italy), at the Austrian SAXS beamline (Amenitsch et al., 1995). Measurements were carried out at 25 °C in the automatic sample changer system developed in the beamline, the µDrop (Haider et al., 2021). This system is configured to measure a minimal quantity of sample, which is placed between two 3 × 1 mm rectangular windows with the observation area made of 2 μm-thick silicon nitride, supported by a 1 mm-wide silicon frame. Three different injections for each sample were carried out to compare measurements. Each SAXS acquisition lasted for 20 s, with a rest time of 2 s for each step, and for each sample injection, 10 acquisitions were obtained. Rest time reduces the possibility of radiation damage. All samples, including buffers, were investigate at the same conditions of temperature and exposure time. Incident and transmitted beam intensities were measured to obtain transmission values. The two-dimensional patterns were recorded with the Pilatus3 1M detector system (Dectris, Switzerland), processed by SAXSDOG (Burian et al., 2022), and by Igor Pro software (WaveMetrics, Lake Oswego, OR, USA) to obtain radial averages. The scattering intensity, expressed as macroscopic differential cross-section dΣ/dΩ, was obtained as a function of the scattering vector Q, defined as Q=4π sinθ/λ, where 2θ is the scattering angle, and λ=0.154 nm is the wavelength of x-rays corresponding to an energy of 8 keV. The sample-detector distance was 1.558 m. The maximum Q was 5.1 nm^-1^. α-crustacyanin in buffer at pH 7.5 was investigated at two concentrations of 1.1 mg/mL and of 0.55 mg/mL. Data analysis was performed using both GENFIT (Spinozzi et al., 2014) and CRYSOL (Franke et al., 2017) software packages, with consistent results.

### In vitro reconstruction and spectrophotometric assay of the HPR-β-crustacyanin unit

The synthetic HPR peptide (sequence EVLNLPSSFHEEGSPISPESATAHVLPTAYQKVDLDS, purity: 96,08%) was purchased from ProteoGenix (Schiltigheim, FR) and used without further purification. The spectrophotometric assays were performed under controlled temperature conditions with a UV-Vis spectrophotometer: Shimadzu UV-1900i (Shimadzu Corporation, Japan). Stepwise additions of the HPR peptide (0.37 mg/ml) to freshly purified β-crustacyanin (0.05 mg/mL) in 50 mM KH₂PO₄ buffer at 5 °C, were performed while monitoring λ*_max_*.

## Data Availability

The crystallographic structure coordinates have been deposited in the Protein Data Bank with the PDB IDs 28QH (β-crustacyanin) and 28TY (α-crustacyanin). The cryo-EM model for α-crustacyanin 12-mer has been deposited with the PDB ID: 28TQ, with Coloumb potential composite map with ID: EMD-56816, consensus map with ID: EMD-56716 and related focused maps with ID: EMD-56713, EMD-56714 and EMD-56715. The cryo-EM model for α-crustacyanin 8-mer has been deposited with PDB ID: 28TZ, with consensus map with ID: EMD-56822. The cryo-EM model for α-crustacyanin 16-mer has been deposited with PDB ID: 28UD, with consensus map with ID: EMD-56823.

## Author Contributions

M.C. initiated, conceived, and supervised the study. A.A., M.C.C., A.D.G. and M.C. designed the experiments. M.C.C., N.R. and A.A. performed protein purification, crystallization, and all functional assays. M.C.C., A.M.C. and C.M.D. performed diffraction data collection. M.C.C. and M.C. solved and refined the crystal structures. M.C.C. and T.W. performed cryo-EM sample preparation and data collection. M.C.C., H.B., A.D.G. and M.C. performed cryo-EM data analysis and reconstructions; M.C.C. and M.C. performed cryo-EM model building and refinement. M.C.C. and M.G.O. performed SAXS data collection and analysis. R.L. and B.D. performed the QM calculations. M.C. and M.C.C. wrote the original draft. All the authors discussed, revised, and approved the manuscript.

## Acknowledgements

X-ray diffraction data were collected at the ESRF (Grenoble, France) storage ring under the beam time award number MX-2464. We would like to thank the staff of the ESRF and EMBL Grenoble for their assistance and support. We are also thankful to the CUNY Advance Science Research Center, New York, NY, US for granting access to the Imaging Facilities. We thank the Proteomics Core Facility at European Molecular Biology Laboratory, (EMBL) Heidelberg (DE) for the service provided. This project has received funding from the European Union’s H2020-MSCA-RISE-2018 Research and Innovation program under the Marie Skłodowska-Curie Grant Agreement No. 823780 (https://prometeus-rise.org/). This research has been partially funded from the project Vitality – Project Code ECS00000041, CUP I33C22001330007 - funded under the National Recovery and Resilience Plan (NRRP), Mission 4 Component 2 Investment 1.5 - ‘Creation and strengthening of innovation ecosystems,’ construction of ‘territorial leaders in R&D’ – Innovation Ecosystems - Project ’Innovation, digitalization and sustainability for the diffused economy in Central Italy – VITALITY’ Call for tender No. 3277 of 30/12/2021, and Concession Decree No. 0001057.23-06-2022 of Italian Ministry of University funded by the European Union – NextGenerationEU. The authors acknowledge the CERIC-ERIC Consortium for the access to experimental facilities and financial support (proposal 20232092) and Dr. Heinz Amenitsch (ELETTRA, Basovizza, IT) for beamline set-up and stimulating discussions. MGO and MC’s research is funded by Ricerca Scientifica di Ateneo (RSA2021–2023) program by the Università Politecnica delle Marche (Ancona, Italy). We thank the Department of Agricultural, Food and Environmental Sciences (D3A) and the Università Politecnica delle Marche (Ancona, Italy) for a Ph.D. studentship award to M.C.C. A.d.G was supported by the US National Institutes of Health (NIH) grant R35 GM133598. MC thanks Prof. John R. Helliwell (University of Manchester, UK), Prof. Paolo Mariani (UNIVPM, Ancona, IT) and Prof. Beatrice Vallone (Università della Sapienza, Rome, IT) for the continuous support.

## Supplementary Materials

Supplementary material for this article is available at http://XXXXX

