## Supplementary material for "Structural basis of the lobster carapace blue colour mediated by an HPR protein": Supplermentary Material

**This PDF file includes:**

Figs. S1 to S5
Tables S1 to S10

**Fig. S1.**

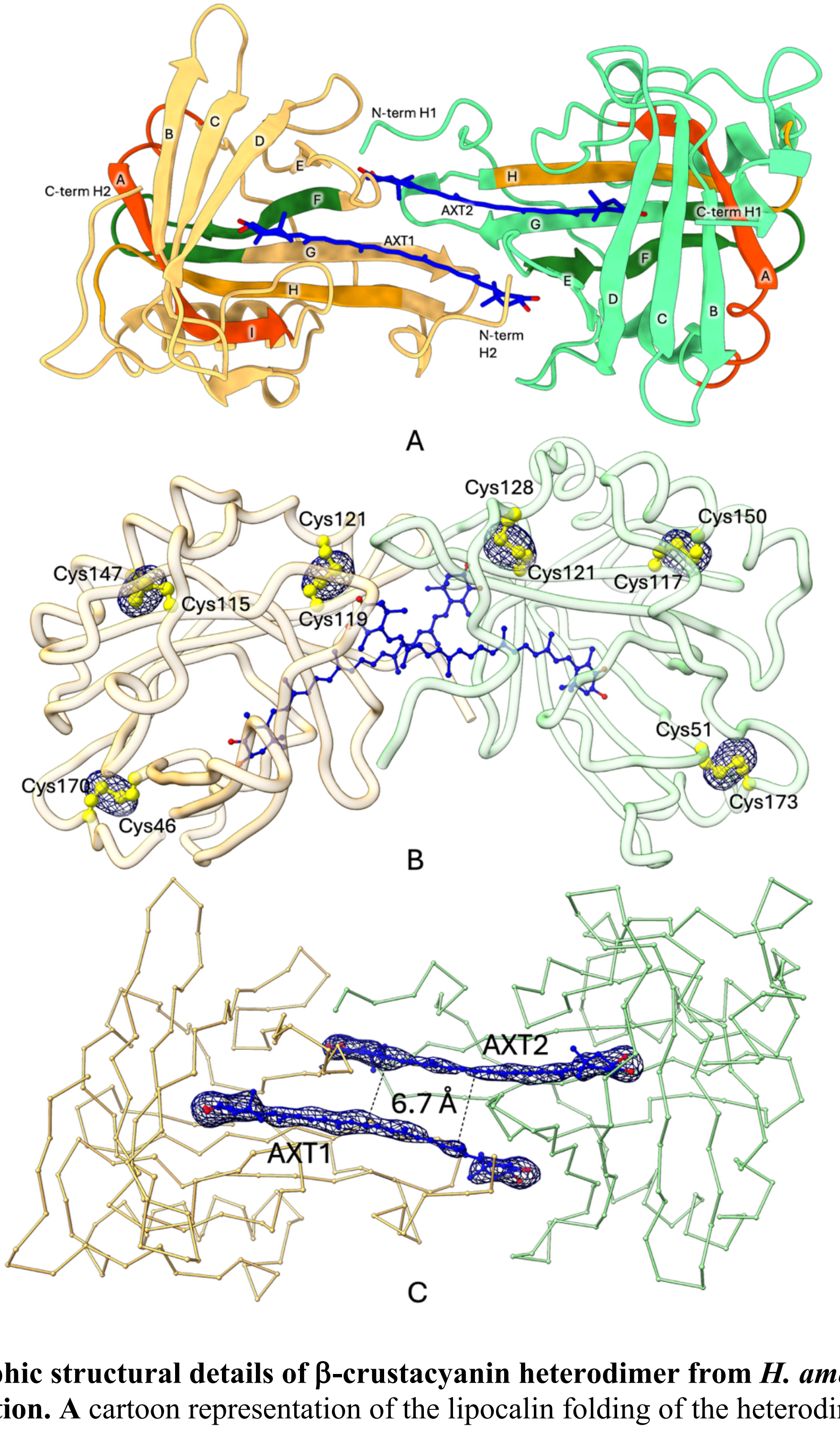


**Crystallographic structural details of β-crustacyanin heterodimer from *H. americanus* at 2.75 Å resolution. A** cartoon representation of the lipocalin folding of the heterodimer (H1, green; H2, beige). **B** Fourier difference omit electron density map (blue colour) countoured at 4σ level around the disulfide bridges depicted in yellow ball-and-stick. **C** Fourier difference omit electron density map (blue colour) countoured at 4σ level around the AXT molecules represented in blue ball-and-stick.

**Fig. S2.**

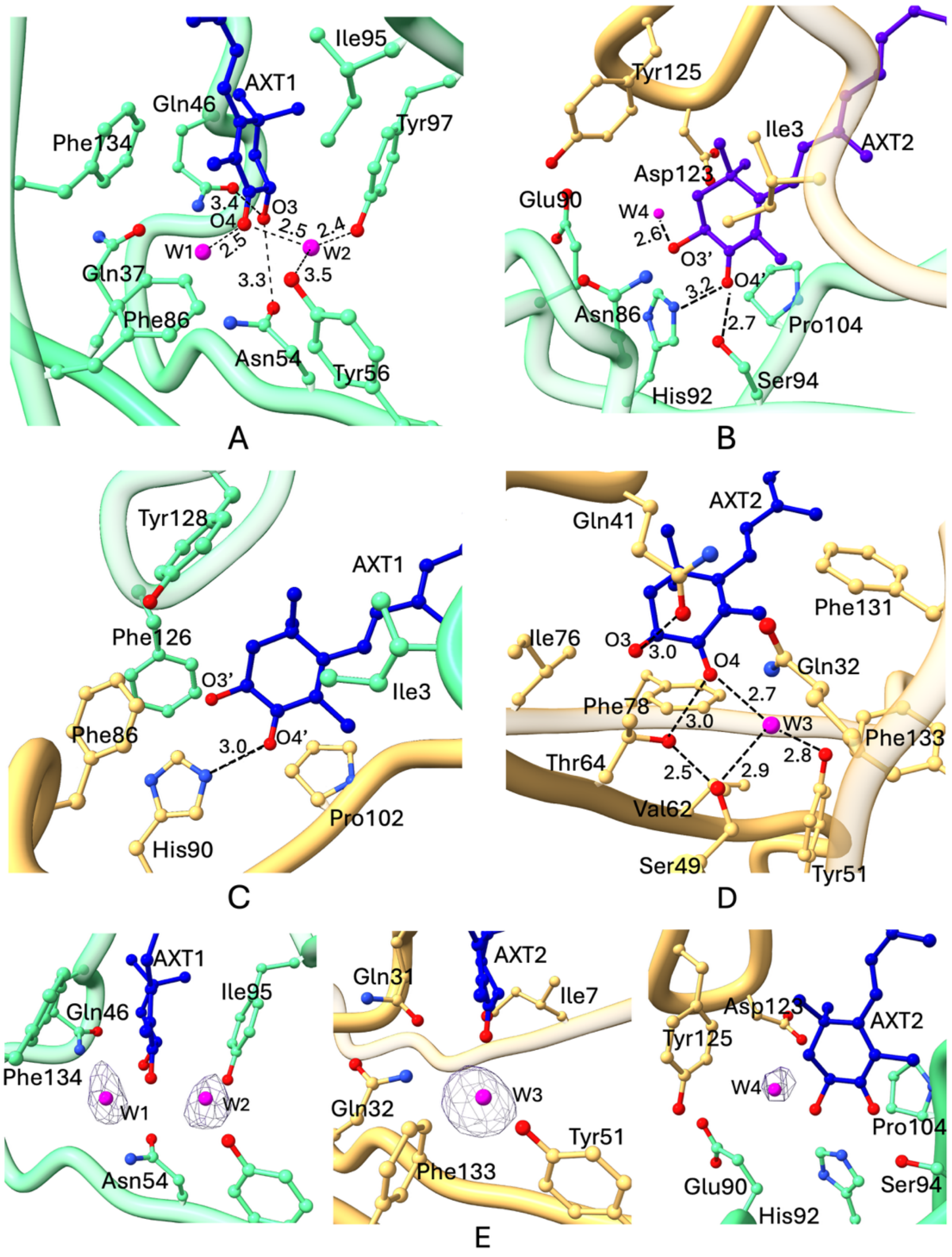


**Crystallographic structural details of *H. americanus* β-crustacyanin AXT binding.** The H1 subunit calyx is coloured in green, the H2 subunit calyx is in beige, and AXT in blue depicted in ball-and-stick. Detailed interactions of the binding site: **A** ring C1-6 AXT1. **B** ring C1’-6’ AXT2. **C** ring C1’-6’ AXT1. **D** ring C1-6 AXT2. **E** Fourier difference omit electron density map countoured at 5σ level (blue colour) around water molecules W1-4 represented in magenta ball-and-stick**.**

**Fig. S3**

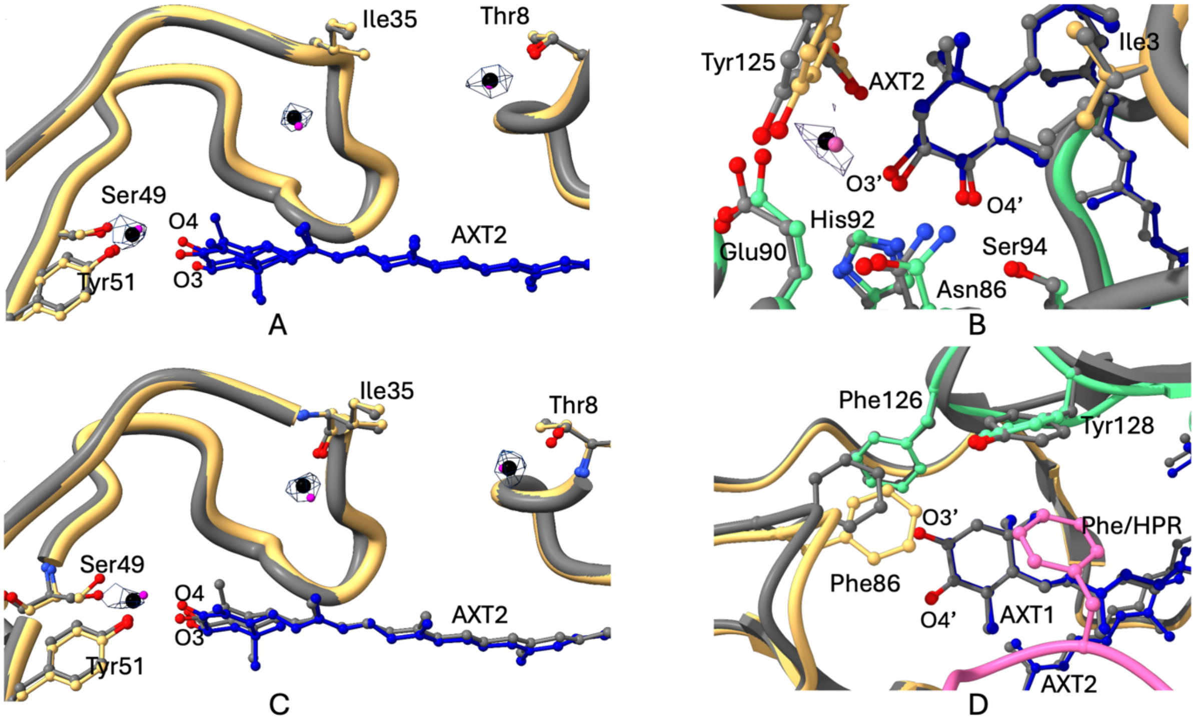


**Superposition of β-crustacyanin H2 subunit determined crystallographically (grey colour) with the β-crustacyanin-HPR complex repeats (beige colour) constituted of monomers**: **a** d; **b** e; **c** f highlighting the waters visible at the C1-6 AXT binding site in the local refined cryo-EM map (blue mesh, contoured at 3.0σ). **d** binding site of AXT β-crustacyanin-HPR complex within H2 subunit showing the proximity of Phe side chain from the HPR chain to the C1'-6' end ring.

**Fig. S4**

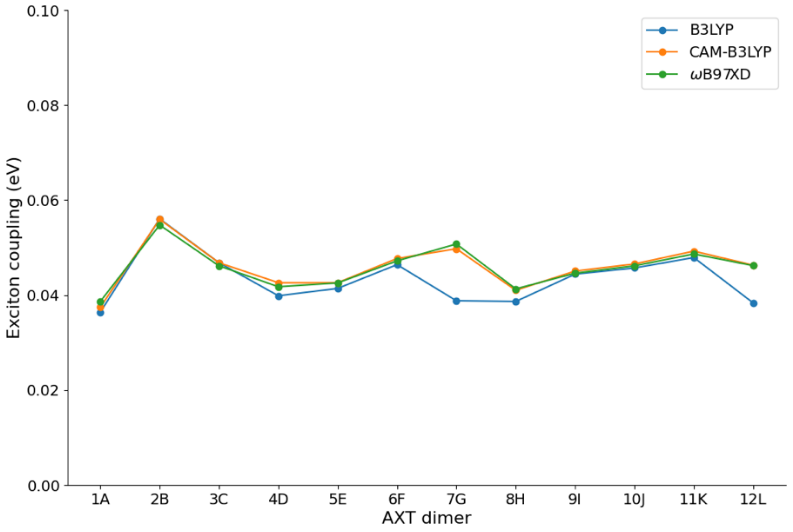


Contribution of exciton coupling to the bathochromic shift calculated for each of the 12 astaxanthin dimers in the cryo-EM structure using different density functionals. The energies used to calculate the exciton coupling are given in Table S10.

**Table S1:** Crystallographic main data collection and refinement statistics for β-crustacyanin and α-crustacyanin from *H. americanus*. Statistics for the highest-resolution shell are given in parentheses. Statistics were calculated using the crystallographic validation tools within PHENIX suite.

| | **β-crustacyanin** | **α-crustacyanin** |
|---|---|---|
| | (PDB code 28QH) | (PDB code 28TY) |
| **Data collection** | | |
| Space group | $P6_322$ | $P222_1$ |
| Cell parameters (a, b, c, Å) | 121.6, 121.6, 184.9 | 123.7, 175.0, 403.4, |
| Resolution range (Å) | 184.9 – 2.75 (2.88 – 2.75) | 132.2 – 6.32 (6.65 – 6.32) |
| **Refinement** | | |
| Reflections used in refinement | 21699 (3017) | 19092 (2628) |
| Reflections used for R-free | 1038 (148) | 1909 (135) |
| Wilson B-factor (Å$^2$) | 79.5 | 459.29 |
| $R_{work}$ [d] | 0.217 (0.377) | 0.2711 (0.3372) |
| $R_{free}$ [d] | 0.258 (0.386) | 0.3736 (0.3939) |
| Total n. of atoms [d] | 2970 | 35283 |
| macromolecules | 2845 | 34315 |
| ligands | 91 | 968 |
| waters | 34 | 0 |
| Protein residues | 355 | 4306 |
| Matthews Coeff (% solvent content) | 4.9 (74.9) | 4.5 (72.9) |
| **r.m.s.d**. [d] | | |
| bond length (Å) | 0.012 | 0.012 |
| angles (°) | 1.89 | 1.58 |
| **Clashscore** | 6.48 | 34.69 |
| **Ramachandran** [d] | | |
| favored (%) | 94.87 | 94.34 |
| allowed (%) | 5.13 | 5.28 |
| outliers (%) | 0.0 | 0.38 |

[a] Values in the highest resolution shell.
[b] $R_{merge} = \sum_{hkl} \sum_j \left| I_j - \langle I \rangle \right| / \sum_{hkl} \sum_j I_j$ , where I is the intensity of a reflection, and $\langle I \rangle$ is

the mean intensity of all symmetry related reflections j.
[c] $R_{p.i.m.} = \sum_{hkl}\left\{[1/(N-1)]^{1/2} \sum_j \left|I_j - \langle I \rangle\right|\right\} / \sum_{hkl} \sum_j I_j$, where I is the intensity of a reflection, and $\langle I \rangle$ is the mean intensity of all symmetry related reflections j, and N is the multiplicity (*90*)
[d] Calculated with PHENIX suite (*32*) $R_{free}$ is calculated using 5% (β-crustacyanin) or 10% (α-crustacyanin) of the total reflections that were randomly selected and excluded from refinement.

**Table S2** Cryo-EM data collection, refinement, and validation statistics. Statistics were calculated using the cryo-EM validation tools within PHENIX suite.

| | (PDB code 28TQ) | (PDB code 28TZ) | (PDB code 28UD) |
|---|---|---|---|
| | Class 0 | Class 1 | Class 0 |
| **Monomeric form** | 12-mer | 8-mer | 16-mer |
| **Data collection and processing** | | | |
| Magnification ( × ) | 37k | | |
| Voltage (kV) | 300 | | |
| Electron exposure ($e^{-}$/Å$^2$/frame) | 1.07 | | |
| Defocus range (μm) (step) | -0.8 to -2.2 (0.2) | | |
| Pixel size (Å) | 0.8465 | | |
| Total number of movies collected (no.) | 10114 | | |
| Final number of particles (no.) | 369648 | 188442 | 181206 |
| Map resolution (Å) | 2.75 | 3.09 | 3.9 |
| **Model Refinement** | | | |
| FSC threshold | 0.143 | | |
| Composition (#) | | | |
| Chains | 51 | 33 | 65 |
| Atoms (Hydrogens) | 36402 (0) | 25665 (0) | 51083 (0) |
| Residues | 4698 | 3132 | 6232 |
| Water | 127 | 0 | 0 |
| Ligands (Astaxanthin) | 24 | 16 | 32 |
| **Refinement** | | | |
| **Bonds (RMSD)** | | | |
| Length (Å) (# > 4sigma) | 0.015 (110) | 0.012 (45) | 0.012 (95) |
| Angles (°) (# > 4sigma) | 2.485 (1018) | 2.311 (460) | 2.444 (1203) |
| MolProbity score | 2.94 | 3.50 | 3.15 |
| Clash score | 21.33 | 37.92 | 25.28 |
| **Ramachandran plot (%)** | | | |
| Outliers | 0.43 | 2.39 | 1.67 |
| Allowed | 5.94 | 11.94 | 8.44 |
| Favored | 93.62 | 85.67 | 89.89 |
| **Rama-Z (Ramachandran plot Z-score, RMSD)** | | | |
| whole | -2.77 (0.11) | -4.19 (0.13) | -3.56 (0.09) |
| helix | -2.56 (0.14) | -3.35 (0.18) | -2.72 (0.14) |
| sheet | -1.14 (0.13) | -1.96 (0.15) | -1.84 (0.11) |
| loop | -2.07 (0.10) | -3.21 (0.12) | -2.62 (0.09) |
| Rotamer outliers (%) | 8.13 | 11.09 | 8.58 |
| Cβ outliers (%) | 9.69 | 4.42 | 5.10 |
| Cis proline/general | 0.0/0.0 | 0.0/0.0 | 0.0/0.0 |

| | | | |
|---|---|---|---|
| Twisted proline/general | 0.0/0.0 | 0.5/0.0 | 0.3/0.1 |
| CaBLAM outliers (%) | 2.48 | 4.96 | 3.43 |
| **ADP (B-factors)** | | | |
| Iso/Aniso (#) | 38623/0 | 25665/0 | 51083/0 |
| min/max/mean | | | |
| Protein | 13.96/640.00/133.01 | 22.30/640.00/261.65 | 45.54/630.94/289.06 |
| Ligand | 18.95/525.23/88.51 | 21.76/609.18/204.69 | 66.15/626.06/272.43 |
| Water | 26.11/87.68/60.78 | | |
| **Occupancy** | | | |
| Mean | 1.00 | 1.00 | 1.00 |
| occ = 1 (%) | 100.0 | 100.00 | 100.00 |
| 0 < occ < 1 (%) | 0.0 | 0.00 | 0.00 |
| occ > 1 (%) | 0.0 | 0.00 | 0.00 |
| **Box** | | | |
| Lengths (Å) | 148.98, 148.14, 302.20 | 144.75, 148.98, 220.94 | 149.83, 156.60, 370.77 |
| Angles (°) | 90.0, 90.0, 90.0 | 90.0, 90.0, 90.0 | 90.0, 90.0, 90.0 |
| Supplied Resolution (Å) | 2.75 | 3.09 | 3.9 |
| Resolution Estimates (Å) | **masked (unmasked)** | **masked (unmasked)** | **masked (unmasked)** |
| d FSC (half maps; 0.143) | ---<br>(---) | ---<br>(---) | ---<br>(---) |
| d 99 (full/half1/half2) | 2.2/---/---<br>(2.0/---/---) | 3.7/---/---<br>(3.6/---/---) | 4.8/---/---<br>(4.5/---/---) |
| d model | 1.8 (1.8) | 3.2(3.2) | 3.9 (3.9) |
| d FSC model (0/0.143/0.5) | 1.7/1.8/2.7<br>(1.7/1.9/3.0) | 3.0/3.1/3.2<br>(3.1/3.1/3.4) | 3.9/3.9/4.5<br>(3.9/4.0/5.5) |
| Map min/max/mean | -0.13/1.81/0.05 | -0.28/0.60/0.01 | -0.10/0.33/0.00 |
| **Model vs. Data** | | | |
| **Refinement type** | **Local** | **Heterogenous** | **Local** |
| CC (mask) | 0.88 | 0.75 | 0.78 |
| CC (box) | 0.85 | 0.80 | 0.88 |
| CC (peaks) | 0.82 | 0.62 | 0.71 |
| CC (volume) | 0.90 | 0.72 | 0.76 |
| Mean CC for ligands | 0.67 | 0.71 | 0.75 |

**Table S3.** Statistics for structure solution with search model polyheterodimers of different length using molecular replacement (Top LLG and TFZ) with Phenix_Phaser (*32*) and followed by rigid body refinement ($R_{factor}$, $R_{free}$) with Phenix_Refine (*32*).

| Model n-mer | Number of solutions | Top-LLG | Top-TFZ | COOT visual inspection | R-factor | R-free |
|---|---|---|---|---|---|---|
| 5 | 5 | 143.77 | 13.6 | Failed | 0.5087 | 0.5419 |
| 6 | 5 | 176.71 | 15.2 | Failed | 0.4879 | 0.5266 |
| 7 | 4 | 196.03 | 16.3 | Failed | 0.4715 | 0.4778 |
| 8 | 5 | 222.9 | 17.8 | Failed | Failed | |
| 9 | 4 | 242.2 | 17.8 | Failed | 0.4585 | 0.5245 |
| 10 | 2 | 209.6 | 17.0 | Failed | 0.4580 | 0.4747 |
| 11 | 2 | 241.8 | 17.3 | Passed | 0.4504 | 0.4783 |
| 12 | 1 | 265.3 | 18.3 | Failed | Failed | |

**Table S4**. Intra-heterodimer interactions between H1 (A, B, C, D, E, F, G, H, I, L and M) and H2 (a,b, c, d, e, f, g, h, i, l and m) subunits, calculated using PDBePISA (www.ebi.ac.uk) (*91*) and visually inspected using COOT (*33*).

| Interactions (H1/H2) | b/B | c/C | d/D | e/E | f/F | g/G | h/H | i/I | l/L | m/M |
|---|---|---|---|---|---|---|---|---|---|---|
| GLY 2 [N] / ASP96 [OD1] | 3.3 | 3.1 | 3.0 | 3.1 | 3.3 | 3.2 | 3.3 | 3.3 | 3.3 | 3.5 |
| TYR 81 [OH] / LYS2 [NZ] | 3.5 | 3.0 | 3.2 | 3.2 | 2.9 | 3.3 | 3.3 | 3.1 | 3.0 | |
| ASP 94 [OD1] / ASP1[N] | 2.6 | 2.8 | 3.4 | 2.7 | 2.7 | 2.7 | 3.2 | 3.3 | 3.4 | 2.7 |
| CYS 119 [O] / ASN125[ND2] | 3.3 | 3.3 | 3.3 | 3.1 | 3.2 | 3.1 | 3.2 | 3.2 | 3.3 | 3.1 |
| THR 121 [N] / ASN 125 [OD1] | 2.9 | 2.9 | 2.9 | 2.9 | 2.9 | 2.9 | 3.0 | 2.8 | 2.9 | 3.0 |
| ASP 123 [OD1] / ASP123[N] | 3.0 | 3.0 | 2.6 | 3.0 | 2.7 | 3.1 | 3.2 | 3.2 | 3.2 | 3.3 |
| ASP 123 [N] / ASP123[O] | 2.9 | 2.9 | 3.0 | 3.0 | 2.9 | 3.0 | 3.0 | 3.1 | 3.0 | 3.1 |
| TYR 125 [OH] / GLU90 [OE1] | 2.8 | 2.6 | 2.6 | 2.8 | 2.5 | 2.5 | 2.5 | 2.5 | 2.5 | 2.7 |

**Table S5**. Interactions between H2 (a, b, c, d, e, f, g, h, i, l and m) subunits of subsequent heterodimers, calculated using PDBePISA (www.ebi.ac.uk) (*91*) and visually inspected using COOT (*33*).

| Interactions (H2/H2) | b/a | c/b | d/c | e/d | f/e | g/f | h/g | i/h | l/i | m/l | n/m |
|---|---|---|---|---|---|---|---|---|---|---|---|
| SER14[OG] / GLU145[OE2] | 2.8 | 2.9 | 2.7 | 3.5 | 2.9 | 3.1 | 3.1 | 3.2 | | 2.7 | 2.6 |
| SER14[O]/PRO142[N] | 3.4 | 3.4 | 3.4 | 3.4 | 3.4 | 3.4 | 3.4 | 3.5 | 3.4 | 3.3 | 3.1 |
| VAL15[O] / GLN138[NE2] | 3.1 | 3.1 | 3.2 | 3.4 | 3.2 | 3.2 | 3.2 | 3.3 | 3.2 | 3.2 | 3.0 |
| ASN17[N] / GLN138[OE1] | 2.9 | 3.0 | 3.2 | 3.0 | 2.9 | 3.4 | 2.9 | 2.9 | 2.8 | 2.7 | 2.6 |
| ASP19[OD1] / TYR31[OH] | 2.4 | 2.4 | 2.5 | 2.5 | 2.4 | 2.4 | 2.5 | 2.5 | 2.3 | 2.4 | 2.1 |
| PHE 86[O] / ALA 168[N] | 2.9 | 2.9 | 3.0 | 3.1 | 2.9 | 2.9 | 3.0 | 3.1 | 2.9 | 3.0 | 3.2 |

**Table S6**. H-bond and salt bridge interactions between HPR repeat (chain X) and H1 (chain A, B, C, D, E, F, G, H, I, L and M) and H2 (chain a, b, c, d, e, f, g, h, i, l and m) subunits of each heterodimer calculated using PDBePISA (www.ebi.ac.uk) (*91*) and visually inspected using COOT (*33*).

| **Interactions (H1/HPR chain)** | **B/X** | **C/X** | **D/X** | **E/X** | **F/X** | **G/X** | **H/X** | **I/X** | **L/X** | **M/X** |
|---|---|---|---|---|---|---|---|---|---|---|
| ASP 1[OD1/2] / SER [OG] (n) | 3.1 | 2.7 | 2.8 | 3.0 | 3.3 | 3.2 | 3.0 | 2.9 | 3.2 | |
| ASP 1[OD1/2] / SER [OG] (n+3) | 3.4 | 3.3 | 3.4 | 3.2 | 3.3 | 3.3 | 3.5 | 3.5 | 3.1 | |
| ILE 3 [O] / HIS[N] | 3.0 | 2.9 | 2.8 | 2.9 | 2.8 | 2.8 | 2.8 | 3.0 | 2.9 | 3.3 |
| ILE 3[N] / SER[O] | 3.1 | 2.8 | 2.8 | 2.9 | 2.8 | 2.9 | 2.8 | 2.9 | 3.0 | 2.8 |
| ASP 5 [OD2] / GLY[N] | 2.8 | 3.1 | 3.4 | 3.0 | 3.0 | 3.2 | 3.2 | 3.2 | 3.1 | 2.8 |
| TYR 45 [0H] / SER[O] | 2.4 | 2.4 | 2.6 | 2.5 | 2.6 | 2.3 | 2.6 | 2.6 | 2.5 | 2.6 |
| SER 100[ O] / ALA [N] | 3.4 | 3.4 | 3.4 | 3.5 | 3.5 | 3.4 | 3.4 | 3.5 | 3.4 | 3.5 |
| SER 100[ O] / THR [OG1] | 3.4 | 3.3 | 3.3 | 2.9 | 3.0 | 3.3 | 2.8 | 3.0 | 3.1 | 3.3 |
| LYS 2 [NZ] / SER[O] | 2.6 | 2.7 | 3.4 | 3.0 | 3.0 | 3.4 | 2.7 | 2.9 | 3.3 | 2.9 |
| **Interactions (H2/HPR chain)** | **b/X** | **c/X** | **d/X** | **e/X** | **f/X** | **g/X** | **h/X** | **i/X** | **l/X** | **m/X** |
| TYR 73[O] / VAL [N] | 2.9 | 2.9 | 3.0 | 3.1 | 2.9 | 3.0 | 3.0 | 2.9 | 2.9 | 3.1 |
| LYS 75[O] / GLN [N] | 2.9 | 2.9 | 2.9 | 3.0 | 2.9 | 3.0 | 3.0 | 2.9 | 3.0 | 3.0 |
| LYS 75[N] / GLN [O] | 3.0 | 3.0 | 3.3 | 3.0 | 3.0 | 3.0 | 3.3 | 3.0 | 3.0 | 3.1 |
| LYS 84[N] / PRO[O] | 3.0 | 3.2 | 3.0 | 3.0 | 2.9 | 3.1 | 2.9 | 2.9 | 2.9 | 3.1 |
| GLU 85[OE] / SER[OG] n | 2.7 | 2.7 | 2.6 | 2.4 | 3.3 | 2.6 | 2.6 | 3.0 | 2.3 | 2.7 |
| GLU 85[OE] / SER [N] n+1 | 2.9 | 3.3 | 3.2 | 3.2 | 3.3 | 3.2 | 3.4 | 3.2 | 3.5 | |
| SER 97[N] / GLU [OE2] | 3.2 | 3.4 | 3.3 | 2.9 | 3.2 | 3.2 | 3.5 | 3.1 | 3.2 | 3.2 |
| GLU 169 [OE] / LEU [N] | 3.2 | 3.4 | 3.2 | 3.5 | 3.3 | 3.2 | 3.5 | 3.3 | | 3.4 |
| ARG 173 [NH1]/ ASP [OD1] | | 2.8 | 3.0 | 3.1 | 3.0 | 3.0 | 3.3 | 3.1 | 3.0 | |

**Table S7**. Interactions between astaxanthin (AXT) molecules and H1 (chain A, B, C, D, E, F, G, H, I, L and M) and H2 (chain a, b, c, d, e, f, g, h, i, l and m) subunits of subsequent heterodimers, calculated using PDBePISA (www.ebi.ac.uk) (*91*) and visually inspected using COOT (*33*). AXT1 is defined as the astaxanthin molecule with ring C1-6 within the cavity of H1 subunit, and AXT2 is defined as the astaxanthin molecule with ring C1-6 within the cavity of H2 subunit.

| **Interactions H1/AXT1** | **B** | **C** | **D** | **E** | **F** | **G** | **H** | **I** | **L** | **M** |
|---|---|---|---|---|---|---|---|---|---|---|
| GLN 46[NE2] / AXT[O3] | 3.1 | 3.0 | 3.0 | 2.8 | 2.5 | 2.6 | 2.6 | 2.8 | 3.2 | 2.8 |
| ASN 54[OD 1] / AXT[O3] | 3.4 | 3.2 | 3.3 | 3.3 | 3.9 | 3.4 | 3.5 | 3.4 | 3.3 | 3.1 |
| **Interactions H2/AXT(H1)** | **b** | **c** | **d** | **e** | **f** | **g** | **h** | **i** | **l** | **m** |
| HIS 90 [NE2] / AXT [O4'] | 3.4 | 2.9 | 3.3 | 3.2 | 3.1 | 3.2 | 3.3 | 3.2 | 3.0 | 3.2 |
| **Interactions H1/AXT2** | **B** | **C** | **D** | **E** | **F** | **G** | **H** | **I** | **L** | **M** |
| ASN 86 [ND2] / AXT [O4'] | 3.1 | 3.2 | 3.2 | 3.2 | 3.2 | 3.2 | 3.1 | 3.2 | 3.2 | 2.7 |
| HIS 92 [NE2] / AXT [O4'] | 3.3 | 3.1 | 3.0 | 2.8 | 3.0 | 3.0 | 2.7 | 2.9 | 2.8 | 2.5 |
| SER 94[OG] / AXT [O4'] | 2.8 | 2.8 | 3.0 | 2.9 | 2.8 | 2.7 | 2.9 | 2.9 | 3.0 | 2.7 |
| **Interaction H2/AXT2** | **b** | **c** | **d** | **e** | **f** | **g** | **h** | **i** | **l** | **m** |
| GLN41 [OE1] / AXT[O3] | 3.4 | 3.5 | 3.2 | 3.3 | 3.5 | 3.6 | 3.3 | 3.0 | 2.9 | 3.2 |
| THR 64[OG1] / AXT[O4] | 2.9 | 3.1 | 3.0 | 3.2 | 2.7 | 2.9 | 2.6 | 3.0 | 2.8 | 2.9 |
| **Interaction H2O/H2** | **b** | **c** | **d** | **e** | **f** | **g** | **h** | **i** | **l** | **m** |
| H2O / SER 49 [OG] | | 3.0 | | 2.6 | 2.4 | 2.6 | | | | |
| H2O / TYR 51[OH] | | 2.6 | | 2.8 | 3.1 | 2.9 | | | | |
| **Interaction H2O/AXT2** | **AXT (b)** | **AXT (c)** | **AXT (d)** | **AXT (e)** | **AXT (f)** | **AXT (g)** | **AXT (h)** | **AXT (i)** | **AXT (l)** | **AXT (m)** |
| H2O / AXT[O4] | | 3.0 | | 3.0 | 2.8 | 3.0 | | | | |
| H2O/AXT[O3'] | | | 2.5 | 2.7 | 3.1 | 2.5 | | | | |
| **Interaction X/AXT2** | **AXT (b)** | **AXT (c)** | **AXT (d)** | **AXT (e)** | **AXT (f)** | **AXT (g)** | **AXT (h)** | **AXT (i)** | **AXT (l)** | **AXT (m)** |
| TYR (n+37)[OH]/AXT[O3'] | 2.4 | 2.6 | 2.4 | 2.5 | 2.5 | 2.4 | 2.3 | 2.7 | 2.6 | 2.5 |

**Table S8.** Structural alignment between astaxanthin (AXT) molecules of H1 and H2 subunits of β-crustacyanin crystallographically determined and astaxanthin (AXT) molecules of H1 (chain A, B, C, D, E, F, G, H, I, L and M) and of H2 (chain a, b, c, d, e, f, g, h, i, l and m) subunits of α-crustacyanin determined using cryo-EM. Alignment r.m.s.d. values (Å) were calculated using ChimeraX (*32*)

| **β-CR AXT** | **α-CR AXT** | | | | | | | | | | | |
|---|---|---|---|---|---|---|---|---|---|---|---|---|
| | **A** | **B** | **C** | **D** | **E** | **F** | **G** | **H** | **I** | **L** | **M** | **N** |
| **H1** | 0.98 | 0.46 | 0.33 | 0.37 | 0.36 | 0.45 | 0.38 | 0.5 | 0.35 | 0.35 | 0.35 | 0.45 |
| | **a** | **b** | **c** | **d** | **e** | **f** | **g** | **h** | **i** | **l** | **m** | **n** |
| **H2** | 0.44 | 0.41 | 0.48 | 0.49 | 0.43 | 0.46 | 0.46 | 0.52 | 0.44 | 0.49 | 0.49 | 0.48 |

**Table S9.** Structural alignment between astaxanthin (AXT) molecules of H1 (chain A, B, C, D, E, F, G, H, I, L and M) and H2 (chain a, b, c, d, e, f, g, h, i, l and m) subunits of α-crustacyanin. Alignment r.m.s.d. values (Å) were calculated using ChimeraX (*32*).

| | **b** | **c** | **d** | **e** | **f** | **g** | **h** | **i** | **l** | **m** | **n** |
|---|---|---|---|---|---|---|---|---|---|---|---|
| **a** | 0.57 | 0.68 | 0.68 | 0.61 | 0.63 | 0.66 | 0.68 | 0.66 | 0.67 | 0.65 | 0.43 |
| **b** | | 0.18 | 0.20 | 0.19 | 0.17 | 0.20 | 0.22 | 0.25 | 0.19 | 0.22 | 0.37 |
| **c** | | | 0.19 | 0.18 | 0.15 | 0.13 | 0.21 | 0.24 | 0.19 | 0.26 | 0.47 |
| **d** | | | | 0.19 | 0.21 | 0.24 | 0.21 | 0.20 | 0.12 | 0.15 | 0.45 |
| **e** | | | | | 0.19 | 0.20 | 0.25 | 0.21 | 0.20 | 0.26 | 0.41 |
| **f** | | | | | | 0.13 | 0.16 | 0.28 | 0.21 | 0.26 | 0.40 |
| **g** | | | | | | | 0.19 | 0.26 | 0.22 | 0.29 | 0.46 |
| **h** | | | | | | | | 0.30 | 0.22 | 0.25 | 0.40 |
| **i** | | | | | | | | | 0.19 | 0.23 | 0.48 |
| **l** | | | | | | | | | | 0.16 | 0.45 |
| **m** | | | | | | | | | | | 0.42 |

| | **B** | **C** | **D** | **E** | **F** | **G** | **H** | **I** | **L** | **M** | **N** |
|---|---|---|---|---|---|---|---|---|---|---|---|
| **A** | 1.14 | 1.03 | 1.02 | 1.08 | 1.09 | 1.06 | 1.09 | 1.06 | 1.05 | 1.08 | 1.06 |
| **B** | | 0.32 | 0.33 | 0.27 | 0.36 | 0.34 | 0.43 | 0.31 | 0.34 | 0.32 | 0.48 |
| **C** | | | 0.16 | 0.22 | 0.26 | 0.17 | 0.33 | 0.13 | 0.18 | 0.25 | 0.38 |
| **D** | | | | 0.18 | 0.29 | 0.16 | 0.29 | 0.16 | 0.24 | 0.25 | 0.40 |
| **E** | | | | | 0.29 | 0.19 | 0.33 | 0.19 | 0.22 | 0.20 | 0.38 |
| **F** | | | | | | 0.21 | 0.28 | 0.27 | 0.30 | 0.34 | 0.45 |
| **G** | | | | | | | 0.23 | 0.18 | 0.21 | 0.23 | 0.40 |
| **H** | | | | | | | | 0.32 | 0.38 | 0.36 | 0.47 |
| **I** | | | | | | | | | 0.20 | 0.21 | 0.38 |
| **L** | | | | | | | | | | 0.25 | 0.40 |
| **M** | | | | | | | | | | | 037 |

**Table S10.** Calculated energies (E) for the two excited states $\Psi_+^*$ and $\Psi_-^*$ of each of the 12 AXT dimers in the cryo-EM structure into which the strongly absorbing monomeric excited state splits as a result of exciton coupling (eV).[a]

| | **ωB97X-D** | | | **B3LYP** | | | **CAM-B3LYP** | | |
|---|---|---|---|---|---|---|---|---|---|
| **Dimer** | $\boldsymbol{E(\Psi_+^*)}$ | $\boldsymbol{E(\Psi_-^*)}$ | **Exciton coupling** | $\boldsymbol{E(\Psi_+^*)}$ | $\boldsymbol{E(\Psi_-^*)}$ | **Exciton coupling** | $\boldsymbol{E(\Psi_+^*)}$ | $\boldsymbol{E(\Psi_-^*)}$ | **Exciton coupling** |
| 1A | 3.051 | 2.974 | 0.039 | 2.480 | 2.408 | 0.036 | 2.996 | 2.921 | 0.037 |
| 2B | 2.932 | 2.822 | 0.055 | 2.361 | 2.249 | 0.056 | 2.873 | 2.761 | 0.056 |
| 3C | 2.976 | 2.884 | 0.046 | 2.393 | 2.300 | 0.047 | 2.917 | 2.824 | 0.047 |
| 4D | 2.984 | 2.900 | 0.042 | 2.393 | 2.313 | 0.040 | 2.874 | 2.788 | 0.043 |
| 5E | 2.934 | 2.849 | 0.043 | 2.353 | 2.270 | 0.041 | 2.874 | 2.788 | 0.043 |
| 6F | 2.954 | 2.859 | 0.047 | 2.376 | 2.283 | 0.046 | 2.894 | 2.799 | 0.048 |
| 7G | 2.947 | 2.846 | 0.051 | 2.328 | 2.250 | 0.039 | 2.885 | 2.785 | 0.050 |
| 8H | 2.931 | 2.849 | 0.041 | 2.341 | 2.264 | 0.039 | 2.870 | 2.788 | 0.041 |
| 9I | 2.979 | 2.890 | 0.045 | 2.390 | 2.301 | 0.044 | 2.920 | 2.830 | 0.045 |
| 10J | 2.957 | 2.865 | 0.046 | 2.367 | 2.275 | 0.046 | 2.897 | 2.804 | 0.047 |
| 11K | 2.960 | 2.863 | 0.049 | 2.382 | 2.286 | 0.048 | 2.902 | 2.803 | 0.049 |
| 12L | 2.938 | 2.845 | 0.046 | 2.344 | 2.267 | 0.038 | 2.878 | 2.786 | 0.046 |

[a] All calculations carried out with the 6-31G(d,p) basis set and using ωB97X-D/6-31G(d,p) geometries of the different dimers.
